# Timely neurogenesis enables increased nuclear packing order during neuronal lamination

**DOI:** 10.1101/2024.11.12.623216

**Authors:** Lucrezia C. Ferme, Allyson Q. Ryan, Robert Haase, Carl D. Modes, Caren Norden

## Abstract

The coordination of cell proliferation, migration, and differentiation is crucial for organogenesis in many tissues, including the central nervous system and other organs that arise from pseudostratified epithelia (PSE). PSE feature densely packed elongated epithelial cells, with nuclei positioned along the apicobasal cell axis in a cell cycle-dependent manner. Also, PSE serve as organ precursors in diverse developmental contexts across evolution. While the role of nuclear movements in PSE has been extensively studied, less is known about whether and how their nuclear packing arrangements and changes of packing state influence tissue morphogenesis. To address this, we analyzed nuclear shapes, sizes and neighborhood statistics by segmenting nuclei in 3D and over development in zebrafish retinal neuroepithelia (RNE). We find that in PSE nuclei exhibit orientational, nematic-like order but remain positionally disordered. This pattern is conserved in other, less packed, neuroepithelia, like the hindbrain, suggesting that nematic-like order is a hallmark of pseudostratification. Our analysis during retinal development also revealed that nuclear packing density increases, approaching theoretical packing limits for disordered monodisperse ellipsoids at stages when the tissue transitions to a laminated neuronal structure. As neurogenesis progresses, nuclear shapes are remodeled, enabling the RNE to shift to a crystalline, ordered structure, while maintaining orientational alignment. Failure to initiate neurogenesis results in severe tissue deformations due to increased buckling instability. Our results thus show an instance where nuclear shape and nuclear positioning and their changes are essential for proper retinal morphogenesis, a phenomenon most likely also found in other tissue arrangements.

## Introduction

Cell proliferation, migration and differentiation need to be tightly coordinated in space and time as well as across scales to ensure that functional organs form reproducibly during development. This coordination is essential in diverse contexts, including the formation of the vertebrate central nervous system (CNS), consisting of spinal cord, retina and brain. Interestingly, all CNS structures arise from pseudostratified epithelia (PSE) [1–7], as do brain and retinal organoids [8–10]. PSE also serve as organ precursors in diverse other developmental contexts both in invertebrates and vertebrates [11–13], like in the organogenesis of lungs and liver in mouse [14, 15] or in the wing imaginal discs in Drosophila or in the embryonic ectoderm in Nematostella [11]. This makes PSE highly conserved organ precursors across evolution. In general, PSE are highly proliferative tissues featuring a monolayer of apicobasally elongated cells that are attached to a basal lamina [13, 16]. Nuclei within these epithelial cells occupy positions all along the apicobasal cell axis during most of the cell cycle, except for during mitosis, which always occurs apically [17–20]. Further, PSE tend to proliferate at high rate during development and are therefore densely packed with nuclei, with densities typically increasing with ongoing proliferation [1, 21]. Together with the broad occurrence of PSE in diverse developmental contexts, the conservation of these features suggests that pseudostratification can aid proper organogenesis. However, how exactly pseudostratification could positively influence neural and other tissue development is not yet fully understood This knowledge gap arises from the fact that so far studies have mainly focused on characterizing cell shape changes and nuclear movements in PSE and how these events affect development [22–27]. However, dense nuclear packing is arguably one of the most distinguishing features of PSE and, despite this, its effects on development have not been well explored. Less is known regarding the effects that dense packing of nuclei might have on morphogenesis of proliferating PSE and whether pseudostratification could play a role in further tissue development. Less is known regarding the effects that dense packing of nuclei might have on morphogenesis of proliferating PSE and whether pseudostratification could play a role in further tissue development. Based on the fact that nuclei occupy a significant fraction of the cell volume and considering that nuclear mechanical properties vary with cell type [28–31], recent studies proposed that nuclear positioning and packing could drive mechanical changes in neuroepithelia of the vertebrate retina and inner ear [32, 33]. Theoretical modelling and analysis of nuclear packing in the densely populated zebrafish retina showed that cells become more constrained and progressively more ordered when nuclear-to-cytoplasmic ratios decrease and nuclear volume fractions approximate limiting packing fractions [33]. While being insightful, these studies were limited to two-dimensional (2D) analyses to describe epithelial arrangements. A 3D appreciation, however, was still lacking due to the considerable technical challenge of producing accurate segmentation and analysis of cells and nuclei in volumetric imaging datasets of crowded tissues. It was recently shown that epithelial cells in PSE feature complex 3D shapes and numerous neighbor exchanges along their apicobasal axis [15, 26]. Thus, an overall in-depth 3D characterization of nuclear packing in neuroepithelia and other PSE is important to capture the complexity of these tissues and provide further insights about the effects that pseudostratification and dense nuclear packing can have for organogenesis.

Here, we addressed these technical challenges using StarDist-3D [34] and quantified nuclear packing densities within proliferating neuroepithelia. We first focused on the developing zebrafish retina and analyzed the transition from PSE to ordered laminated neuronal structure. During retinal development, multipotent progenitor cells in the PSE proliferate to sustain tissue growth while concomitantly starting to generate neurons [21]. These neurons, upon birth at the apical surface, relocate within the retina to the positions at which they later function [35, 36]. The emergence of neuronal layers ensures proper connectivity and functionality of the neural tissue [37]. When this spatial ordering of the neurons is impaired, mature tissues can become pathological and dysfunctional [38, 39]. While many studies have shed light on how the intracellular mechanisms of proliferation, differentiation and migration are genetically encoded and regulated [36, 40, 41], less is known about the influence that nuclear packing changes could have on the morphogenesis of the retina. It is also not clear to what extent nuclei are arranged in an ordered or disordered fashion in the retinal PSE and how further ordered tissue structures physically arise during the development from the less ordered PSE into a layered neuronal architecture.

Our systematic 3D analysis showed that nuclear volume fractions increased continuously over the proliferative phase, reaching theoretical packing fraction limits upon neuronal lamination. We find that nuclei are orientationally ordered and positionally disordered in the proliferating RNE, reminiscent of the nematic-like order prevalent in liquid crystals. This orientational order is conserved in a differently shaped neuroepithelium that shows looser packing regimes, the developing hindbrain. Together, this indicates that nematic-like order is a hallmark of pseudostratification. Later in development, neurogenesis progressively remodels nuclear shapes and aids in the transition from the RNE to an even more positionally ordered, crystal-like structure, while maintaining the overall orientational order of nuclei in the laminated retina. Together, our results suggest that progressive emergence of nuclear packing order in the RNE enables the close arrangement of newly formed neurons. When this is impaired, tissue shape is not maintained and further development is perturbed.

## Results

### Instance 3D segmentation of nuclei in the zebrafish retinal neuroepithelium

To analyze the arrangements of progenitor cell nuclei within the retinal neuroepithelium (RNE) over development, we segmented single nuclei using the tg(hsp70:H2B-RFP) zebrafish transgenic line, which marks chromatin and labels all nuclei. The RFP labelling allowed us to image with lasers at longer wavelengths reducing light scattering and improving penetration depth. Furthermore, the variable expression of H2B-RFP under a heat-shock promoter facilitated more efficient segmentation compared to more evenly distributed staining, like DRAQ5, or constructs expressed under a constitutive promoter (Fig. S1 A, B). Tg(hsp70:H2B-RFP) embryos were staged and fixed starting at 24 hours post fertilization (hpf), when the optic cup consists of only proliferating progenitors, every 6 ± 1 hours until 48 hpf, by which time neuronal lamination is ongoing (Fig. 1 A). Locating and separating individual nuclei within a 3D volume is a computer vision task, called instance segmentation, which assigns a label mask to each single object. We annotated 14 cropped 3D images with approximately 40 to 300 labelled nuclei each and used this dataset to train a StarDist-3D model [34], which we then applied for segmentation in the volumetric dataset (Fig. S1 C, D; Table 4). Based on this dataset, we established an image analysis pipeline to extract several nuclear shape descriptors, such as nuclear size and axes’ length, and explore the arrangement of nuclei within the RNE over time (Fig. S2 & S3). For further details on the exact analysis pipeline, see the Methods section and the schematic in Fig. S2 A.

**Fig. 1.**
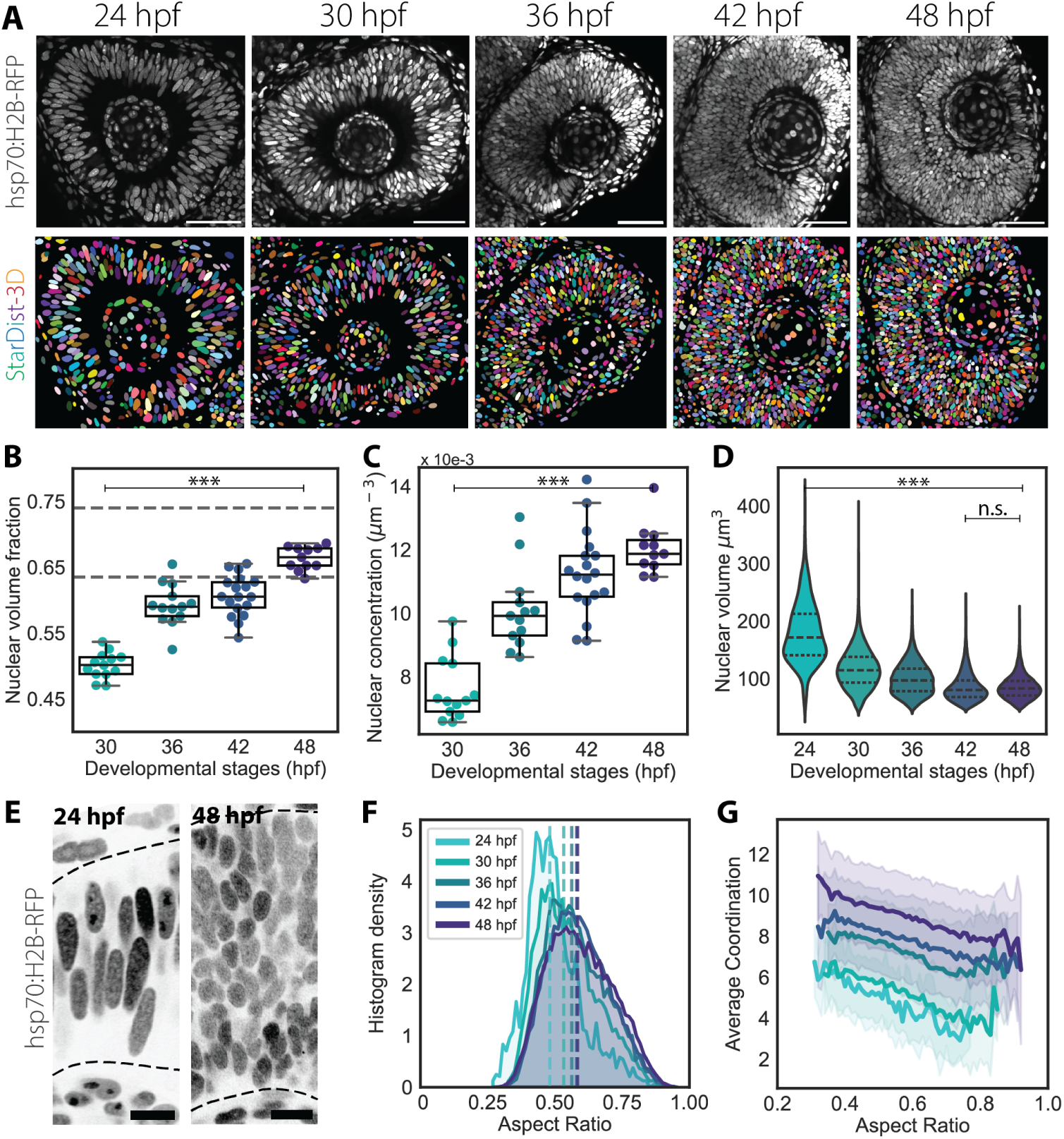
Instance 3D segmentation of nuclei shows increasing nuclear packing densities in the retinal PSE. **A)** Representative confocal sections of the developing retina between 24 hpf and 48 hpf (top row) and corresponding segmented predictions obtained using StarDist-3D (bottom row). Nuclei are labeled with tg(hsp70:H2B-RFP). Scale bar is 50 *µ*m. **B)** Volume fraction occupied by nuclei in regions of interests (ROIs). Segmented lines show the volume fractions *ϕ_s_* = 0.635 and *ϕ_e_* = 0.74 for RCP densities of isovolumetric spheres and fully aspherical ellipsoids, respectively. 30 hpf, N = 13 embryos; 36 hpf, N = 13; 42 hpf, N = 18; 48 hpf, N = 11. **C)** Concentration of nuclei found within the ROIs of panel (B). P values for alpha = 0.01: *** *<* 0.0001 from unpaired two-tailed Student’s t test. D) Distributions of nuclear volumes. P values for alpha = 0.01: *** *<* 0.0001; n.s. not significant; results from Mann Whitney U test. E) Representative confocal sections of the 24 and 48 hpf retina from panel (A). Segmented lines mark the apical and basal boundaries of the retina. Scale bar is 10 *µ*m. **F)** Histogram density distributions of nuclear aspect ratios, which were defined as the ratio between the longer axis length and the mean length between the intermediate and shorter axis. **G)** Correlation between mean number of contacts (average coordination) and nuclear aspect ratios. Solid lines show mean values, shaded area indicate the standard deviation from the mean.

### Nuclear packing in the retinal PSE reaches theoretical close packing densities over development while nuclei keep an ellipsoidal shape

To evaluate nuclear packing, we quantified nuclear counts and the volume fraction they occupy within defined regions of interests (ROI) in the central part of the RNE (see Methods). We found that the nuclear concentration, i.e. number of nuclei per volume, and the nuclear volume fractions increased linearly between 30 and 48 hpf (Fig. 1 B, C). This increase was accompanied by a progressive decrease of nuclear volumes between 24 and 42 hpf, until the onset of neuronal lamination (Fig. 1 D). Interestingly, the median volume fraction *ϕ* reached values up to *ϕ_s_* ≈ 0.635 (Fig. 1 B), a value that is generally known as the Random Close Packing (RCP) density for the disordered packing of isovolumetric spheres reaching their limiting maximum density [42]. The RCP density varies greatly between 2D and 3D situations, so that the RCP value *ϕ_d_* for monodisperse discs in 2D is higher than for spheres in 3D and falls in the range 0.84*<ϕ_d_<*0.89 [43], showing the relevance of considering nuclear packing in 3D. Note that the RCP density describes the densest possible packing of objects without any short- or long-range order, so that particles in the system can no longer move freely and are “locked” in place by their closest neighbors [44]. However, nuclei in the RNE are not simply isovolumetric spheres but more closely resemble ellipsoids (Fig. 1 E). Ellipsoidal particles are known to be able to pack more densely than *ϕ_s_*, reaching RCP densities in the range 0.64*<ϕ_e_<*0.74 [45, 46]. By 48 hpf, most of the analyzed retinas had volume fraction values higher than *ϕ_s_* and in the range of *ϕ_e_*, implying that the retinal PSE was possibly approximating a jamming transition boundary for ellipsoidal particles. To test how nuclear shapes responded to increased packing, we quantified nuclear shapes and monitored how they changed as packing increased. Our analysis showed that most nuclei retained their average ellipsoidal shape throughout the proliferative phase and that nuclear volume reduction resulted mainly from a progressivee shortening of the major axis until 42 hpf (Fig. 1 F; Fig. S4 A,E,F; Table 1). In contrast, volume distributions of nuclei between 42 hpf and 48 hpf, which corresponds to the onset of neuronal lamination, did not change (Fig. 1 D; Table 1), suggesting that nuclear size changes could be affected by nuclear packing and by neuronal lamination.

**Table 1.** Nuclear volume, axes lengths and aspect ratios of nuclei in the central retinal neuroepithelium and in the hindbrain neuroepithelium. Mean and standard deviation are shown for each stage. Mann-Whitney tests between previous and following stage were performed on nuclear volume distributions and aspect ratio distributions, i.e. comparing distributions in the retinal neuroepithelium between 24 hpf and 30 hpf, between 30 hpf and 36 hpf, between 36 hpf and between 42 hpf and 48 hpf. Same procedure was performed for the hindbrain. P-values are shown for each comparison.

| RETINAL NEUROEPITHELIUM |  |  |  |
| --- | --- | --- | --- |
| Nuclear volume($\mu\text{m}^3$) | | | |
| Stage | $\mu \pm \sigma$ | H0: same distribution | p value |
| 24 hpf | 177.87 ± 55.77 | – | – |
| 30 hpf | 113.52 ± 39.92 | (24 hpf) = (30 hpf) | $p \ll 0.0001$ |
| 36 hpf | 95.76 ± 30.54 | (30 hpf) = (36 hpf) | $p \ll 0.0001$ |
| 42 hpf | 81.67 ± 23.40 | (36 hpf) = (42 hpf) | $p \ll 0.0001$ |
| 48 hpf | 81.92 ± 23.06 | (42 hpf) = (48 hpf) | 0.1798 |
| Nuclear axis lengths ($\mu\text{m}$) | | | |
| Stage | A ($\mu \pm \sigma$) | B($\mu \pm \sigma$) | C ($\mu \pm \sigma$) |
| 24 hpf | 11.5 ± 2.59 | 6.17 ± 0.87 | 4.81 ± 0.66 |
| 30 hpf | 9.08 ± 2.00 | 5.50 ± 0.9 | 4.31 ± 0.70 |
| 36 hpf | 8.34 ± 1.68 | 5.25 ± 0.78 | 4.22 ± 0.57 |
| 42 hpf | 7.85 ± 1.43 | 5.00 ± 0.71 | 4.04 ± 0.57 |
| 48 hpf | 7.85 ± 1.33 | 5.02 ± 0.72 | 4.08 ± 0.58 |
| Aspect ratio (A/(B+C)) |  |  |  |
| Stage | $(\mu \pm \sigma)$ | H0: same distribution | p value |
| 24 hpf | 0.49 ± 0.101 | – | – |
| 30 hpf | 0.55 ± 0.111 | (24 hpf) = (30 hpf) | $p \ll 0.0001$ |
| 36 hpf | 0.58 ± 0.113 | (30 hpf) = (36 hpf) | $p \ll 0.0001$ |
| 42 hpf | 0.59 ± 0.111 | (36 hpf) = (42 hpf) | 0.016 |
| 48 hpf | 0.59 ± 0.120 | (42 hpf) = (48 hpf) | 0.899 |

**Table 1** Nuclear volume, axes lengths and aspect ratios of nuclei in the central retinal neuroepithelium and in the hindbrain neuroepithelium.
| HINDBRAIN NEUROEPITHELIUM |  |  |  |
| --- | --- | --- | --- |
| Nuclear volume($\mu\text{m}^3$) | | | |
| Stage | $\mu \pm \sigma$ | H0: same distribution | p value |
| 18 hpf | 315.68 ± 41.88 | – | – |
| 22 hpf | 279.37 ± 46.43 | (18 hpf) = (22 hpf) | 0.2323 |
| 24 hpf | 241.93 ± 75.72 | (22 hpf) = (24 hpf) | $p \ll 0.0001$ |
| 30 hpf | 201.66 ± 20.11 | (24 hpf) = (30 hpf) | $p \ll 0.0001$ |
| Nuclear axis lengths ($\mu\text{m}$) | | | |
| Stage | A ($\mu \pm \sigma$) | B($\mu \pm \sigma$) | C ($\mu \pm \sigma$) |
| 18 hpf | 9.34 ± 1.82 | 6.57 ± 0.77 | 5.63 ± 0.70 |
| 22 hpf | 9.68 ± 1.80 | 6.48 ± 0.77 | 5.45 ± 0.66 |
| 24 hpf | 8.99 ± 1.61 | 6.09 ± 0.68 | 5.01 ± 0.64 |
| 30 hpf | 7.97 ± 1.49 | 5.79 ± 0.81 | 4.87 ± 0.61 |
| Aspect ratio (A/(B+C)) |  |  |  |
| Stage | $(\mu \pm \sigma)$ | H0: same distribution | p value |
| 18 hpf | 0.67 ± 0.13 | – | – |
| 22 hpf | 0.63 ± 0.13 | (18 hpf) = (22 hpf) | $p \ll 0.0001$ |
| 24 hpf | 0.64 ± 0.13 | (22 hpf) = (24 hpf) | 0.292 |
| 30 hpf | 0.69 ± 0.10 | (24 hpf) = (30 hpf) | $p \ll 0.0001$ |

To explore the relation between elongated ellipsoidal morphology and RCP densities, we compared the number of nearby neighbors per nucleus to their aspect ratio. Because neighboring nuclei do not physically touch due to the presence of cell membranes between them, nearby neighbors were defined as those nuclei that are in contact after a small dilation of each nucleus’ outline boundaries, here referred to as touching neighbors (Fig. S3 D, E). After testing different radii of dilation, we adopted a radius of 0.25 *µ*m as minimal effective touching-through-cell-membrane length scale, since this number was the minimal step size of voxels in our imaging dataset. This analysis confirmed that the average contact number Z and the nuclear aspect ratio of each nucleus were linearly correlated, meaning that more elongated nuclei tend to contact a higher number of neighbors, also referred to as the average coordination number (Fig. 1 G). This correlation between average coordination and aspect ratio did not vary across stages, while the range of average coordination increased, reaching contact number values closer to 10 by 48 hpf. The maximum contact numbers for ordered arrangements of isovolumetric spheres and ellipsoids are known to be at *Z_s_*=12 and *Z_e_*=14, respectively [45]. This again shows that nuclei in the RNE approach limiting RCP densities at the onset of neuronal lamination.

Overall, the quantification of nuclear shape descriptors from 3D instance segmentation enabled us to follow nuclear packing changes in the RNE over development. Our results indicate that nuclear shape and size in progenitor cells determine nuclear packing densities in the proliferating RNE and allow the tissue to reach high nuclear volume fractions,without impeding progenitor nuclear movements [47, 48]. Our image analysis code and the trained StarDist-3D model are openly accessible (see Methods) and we are confident that these will be useful for researchers working on similar topics.

### Nuclei are arranged in a nematic-like order in the proliferative phase

We asked whether the increase in nuclear volume fractions in the RNE during the proliferative phase might be explained by the possible emergence of ordered nuclear arrangements. To test this idea, we analyzed whether during the proliferative phase nuclear packing varied spatially within the retinal PSE and/or whether nuclei showed positional order.

One way to evaluate the nuclear packing environment is by looking at the average coordination number of nuclei across the apicobasal axis of the tissue. To do this, we performed image smoothing by averaging our measurements across the voxels of our segmentation dataset and visually inspected the result (Fig. S2 B, C). We found that the average coordination of nuclei in the PSE increased continuously between 24 and 48 hpf, reaching higher values in the inner part of the tissue (Fig. 2 A). The average contact numbers were lower in the retinal ganglion cell layer (RGL), which formed between 42 and 48 hpf, suggesting that neuronal lamination locally reduced nuclear packing densities locally (Fig. 2 A). To discriminate between a scenario in which random nuclear packing is simply increasing and a scenario in which the positional arrangement of nuclei is changing at the same time, we analyzed the mean internuclear centroid-to-centroid distance of touching neighbors. We found that this distance initially decreased, but plateaued at around 5 *µ*m between 30 and 48 hpf, while the mean contact numbers continued to increase (Fig. 2 C). This minimal internuclear distance was confirmed by analysis of proximal neighbors within a set radius, which showed that the number of such neighbors rose between 5 and 10 *µ*m radii (Fig. 2 D). These results reinforced the previous observations that nuclear packing in the developing tissue becomes denser between stages 30 hpf and 48 hpf. Next, we explored whether any periodic positional ordering of nuclei emerged over development by computing the radial distribution function (RDF). This function describes how particle density changes as a function of distance between particles. Consequently, the RDF provides insights on the positional order of particles which are in our case the nuclei. Our results indicated the highest probability to find nuclei at around 5 *µ*m internuclear distance, matching the minimal internuclear distance observed above and thus associated to nearest neighbors, whilst the RDF smoothly decreases at longer distances (Fig. 2 E). Such a profile in the RDF is consistent with a disordered liquid-like arrangement, thereby indicating that between 24 hpf and 48 hpf no regular positional order is present, which would have led to the appearance of more peaks in the RDF. So far, our analysis shows that nuclei pack in a disordered state between 24 and 48 hpf. At the same time, nuclei feature an anisotropic morphology throughout the proliferative phase. Depending on the extent of the alignment, anisotropic nuclei could be arranged in an orientational order in the RNE. This, together with the positional disorder shown by the RDF, would indicate the presence of a collective arrangement of nuclei in a nematic-like orientational order. Nematic order represents the simplest liquid crystalline order, which describes a physical state intermediate to a liquid and a solid, with only partially broken continuous symmetry. Such liquid crystals feature anisotropic objects that are aligned along a common direction axis, even though they lack positional order. To understand if nuclei in RNE were organized in this manner, we calculated the primary nematic axis of each nucleus (Fig. S3 G, H) and their misalignment angle to the average angle of surrounding nuclei from our imaging dataset. We found that nuclei were strongly aligned throughout development, with an average misalignment angle of only around 10 degrees (Fig. 2 B). Computing the local orientation order parameter further confirmed the presence of strong alignment (Fig. 2 F). The same analysis was performed for a secondary nematic axis, which was defined as the elongation axis on the midplane to the primary nematic (Fig. S3 G, I). Also this analysis revealed the presence of a consistent alignment, though weaker than that along the primary axis (Fig. 2 B, F). This secondary alignment suggests that nuclei in the RNE are actually organized like in a biaxial nematic.

**Fig. 2.**
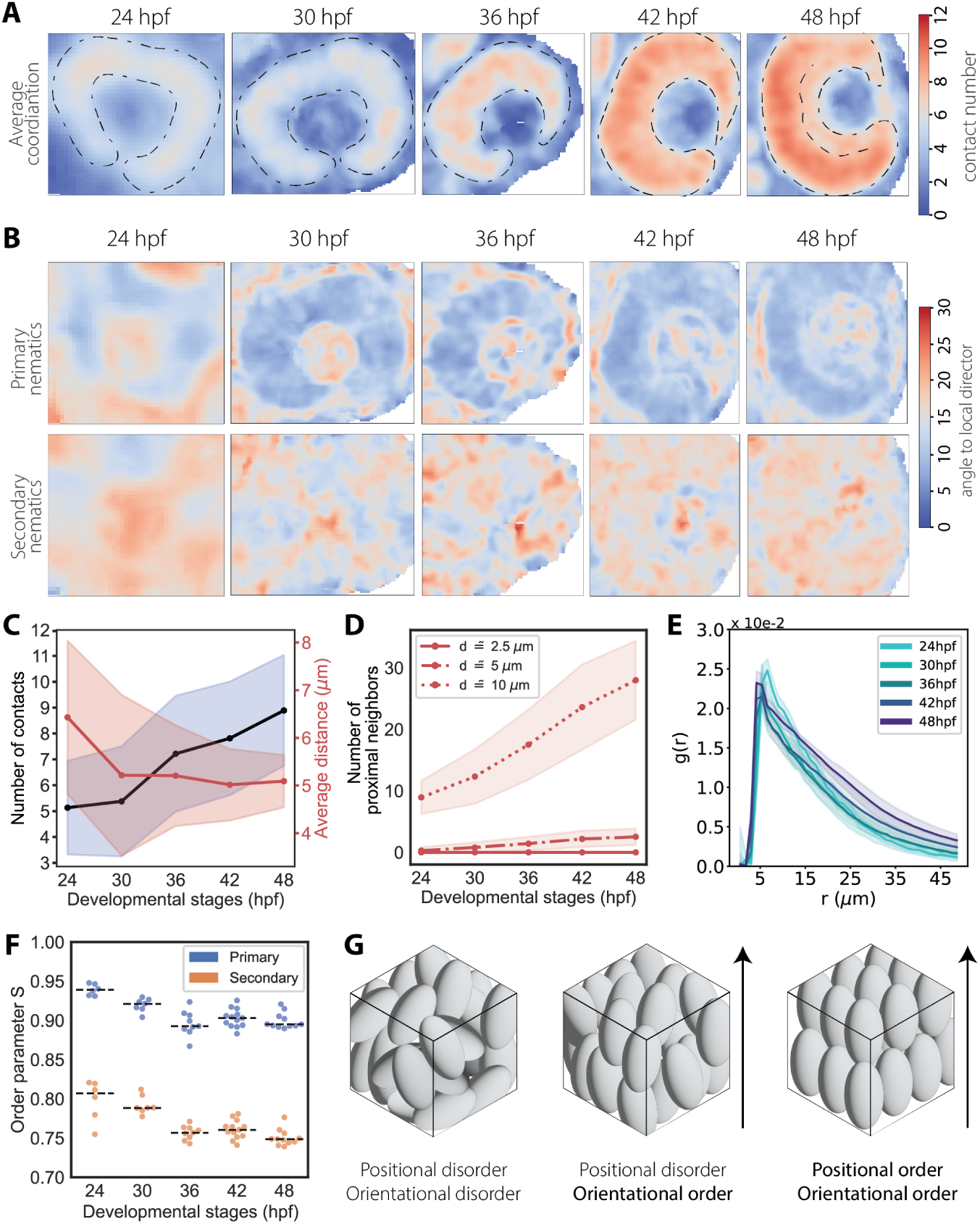
Nuclei are positionally disordered but orientationally ordered in the retinal PSE. **A)** Median intensity projections of representative smoothened imaging datasets reporting the average coordination number for nuclei within a given 3D window of size 10×10×10 *µ*m. The representative images shown are the same as in figure 1 A. Outline of the retinal PSE is marked by segmented lines, together with the space separating the retinal ganglion cell layer from the rest of the tissue at 48 hpf. **B)** Median intensity projection of representative smoothed imaging datasets reporting the mean angle of misalignment between primary nematic axes and the local primary nematic direction (upper row) and the mean angle of misalignment between secondary nematic axes and the local secondary nematic direction (bottom row). Measurements were collected and averaged within a given 3D window of size 10×10×10 *µ*m (upper row). **C)** Mean number of contacts within ROIs increase, while the average distance between touching neighbours stays constant. **D)** Mean number of proximal neighbours per nucleus within different distance radii. **E)** The RDF g(r) calculated for nuclei in ROIs at different developmental stages between 24 and 48 hpf indicate a liquid-like state. In panels (**C, D, E**), solid lines always correspond to mean values, shaded area mark the standard deviation from the mean. **F)** Order parameter S over development. **G)** Examples of monodisperse ellipsoids arranged in different configurations, from fully disordered to fully ordered packing. The arrow represents the average direction of alignment.

Overall, our analysis demonstrated that packing densities vary spatially within the RNE and that nuclei are positionally disordered while being orientationally ordered (Fig. 2 G). This suggests that nuclei in the RNE are arranged with biaxial nematic order, similar to that found in many liquid crystals. This indicates that nuclei in the retinal PSE are positionally disordered and can move through the tissue as in a liquidlike arrangement, but present one degree of order as in a more solid-like configuration.

### Nematic-like order is a hallmark of pseudostratification also at looser packing regimes

It was previously shown that nuclear positioning mechanisms vary between differently shaped PSE within the same organism, such as the developing zebrafish retina and hindbrain [19, 49]. While the RNE is characterized by a hemispherical morphology, the developing hindbrain is a straight tubular PSE. We asked whether despite these differences in architecture and nuclear translocation mechanisms, nuclear packing densities and ordering were conserved or whether they differed between RNE and hindbrain during proliferation and onset of differentiation. To this end, we carried out the same 3D analysis in the developing hindbrain as done in the retinal PSE. Hindbrain neuroepithelia were analysed at set intervals between 18 and 30 hpf, i.e. between the formation of the neural rod and delamination of newly formed neurons [50–52], as shown by HuC/HuD staining (Fig. 3 A, B; Fig. S5 A). In contrast to the RNE, we found that nuclear volume fraction and nuclear concentration did not increase until neuronal lamination at 30 hpf (Fig. 3 C; Fig. S4 B), while still showing a positive correlation like in the retinal PSE (Fig. S5 C). Between 18 and 24 hpf, nuclear packing remained in a looser packing regime than what was found in the RNE, while nuclear volumes decreased over time starting around 24 hpf at the onset of neuronal lamination (Fig. 3 D; Table 1). Even though nuclei presented aspect ratios closer to 1, indicating a more spherical shape compared to nuclei in the RNE, and nuclear volume reduced isotropically (Fig. 3 E; Fig. S5 E, F), nuclear morphologies still inversely correlated with average coordination numbers (Fig. S5 G). The mean internuclear distance changed marginally between 18 hpf and 30 hpf (Fig. S5 H, L), though the average numbers of touching neighbors after dilation and of proximal neighbors within a 10 *µ*m radius significantly increased between 24 hpf and 30 hpf (Fig. 3 F; Fig. S5 I, L). Moreover, nuclei in the hindbrain were still arranged with nematic order as measured by the RDF and the alignment of the primary nematic axes (Fig. 3 G, H, I). Together, this suggests that the arrangement of nuclei with nematic order is conserved across different PSE, regardless of their packing density or tissue shape.

**Fig. 3.**
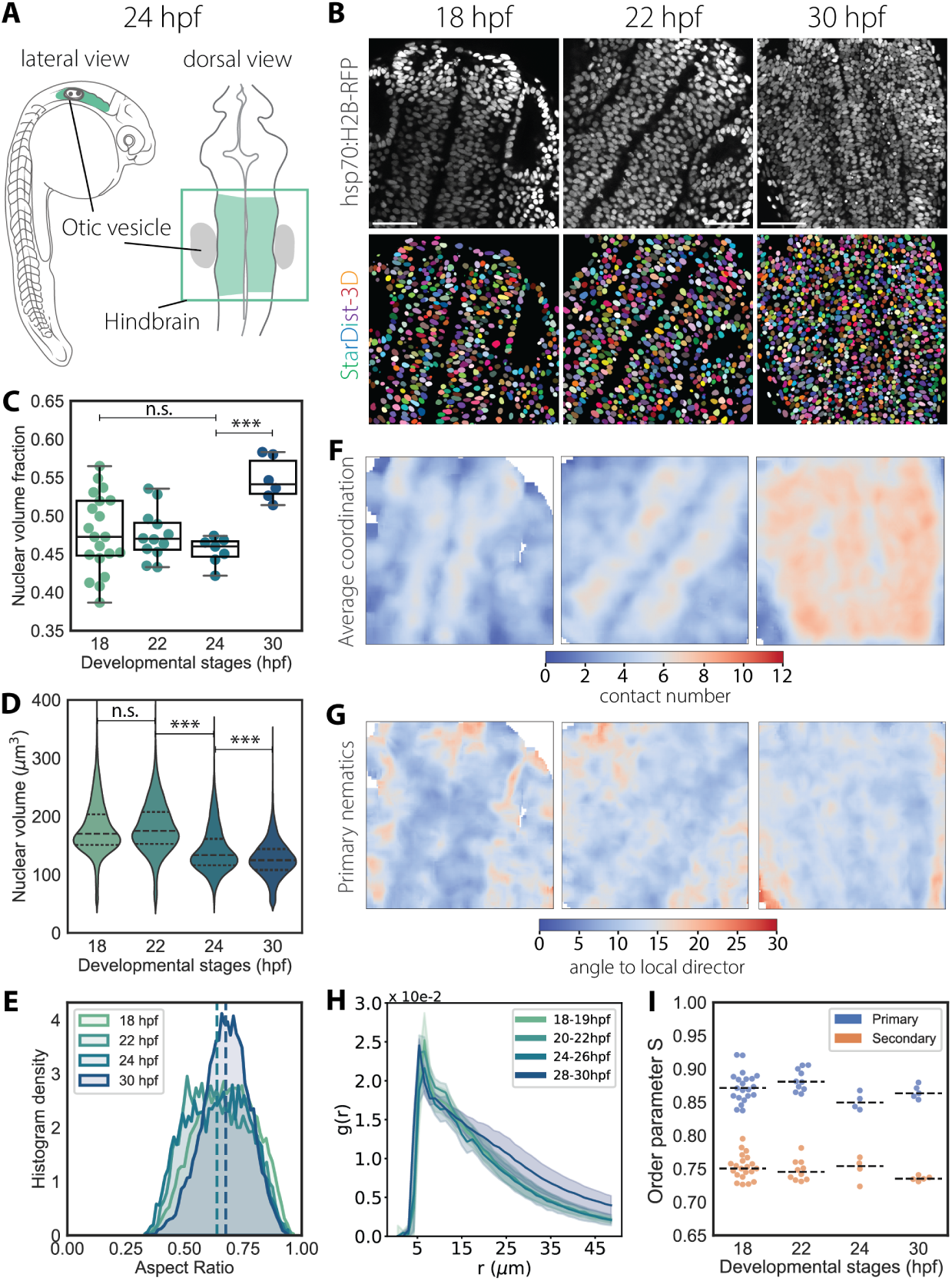
Instance 3D segmentation of hindbrain PSE reveals nematic ordering of nuclei at lower packing regimes. **A)** Scheme of a 24 hpf zebrafish marking the region of the hindbrain that was imaged and quantified. **B)** Representative images of the hindbrain PSE during early development (upper row) and corresponding instance segmentation using StarDist-3D model (bottom row). Nuclei are labelled with Tg(hsp70:H2B-RFP). Scale bar is 50 *µ*m. **C)** Nuclear volume fractions over time. The segmented grey line marks *ϕ_s_* = 0.635 for nuclear volume fraction. 18 hpf, N = 20 embryos; 22 hpf, N = 11; 24 hpf, N = 7; 30 hpf, N = 6. P values for alpha = 0.01: *** *<* 0.0001; n.s. not significant; from unpaired two-tailed Student’s t test. **D)** Violin plot of nuclear volume distributions over time. Segmented lines indicate first and third quartiles and medians. P values for alpha = 0.01: *** *<* 0.0001; n.s. not significant; results from Mann Whitney U test. **E)** Histogram density distributions of nuclear aspect ratios. Segmented lines indicate the mean. **F)** Median intensity projections of representative smoothed imaging datasets reporting the average coordination number for nuclei within a window of size 10×10×10 *µ*m. The representative images shown are the same as in panel (B). **G)** Median intensity projection of representative smoothed imaging datasets reporting the mean angle of misalignment between primary nematic axes and the local primary nematic direction within a window of size 10×10×10 *µ*m. The representative images shown are the same as in panel (B). **H)** The RDF g(r) between 18 and 30 hpf. Solid lines correspond to mean values, shaded area indicate the standard deviation from the mean. **I)** Order parameter S for primary and secondary nematics.

### Neurogenesis coincides with nuclear shape remodeling and aligned crystalline-like arrangements within the laminating retina

Our analysis of nuclear packing in the RNE over development showed that nuclear volume fractions increased during the proliferative phase and reach the theoretical limiting density *ϕ_s_* at the onset of neuronal lamination. In addition, we observed spatial changes in nuclear arrangements upon neurogenesis and during the formation of the RGL. Because cell fate specification entails cell shape changes and repositioning within the tissue, we compared nuclear shape distributions and arrangements between the different neuronal layers: the RGL, the outer nuclear layer (ONL) hosting photoreceptor cells and the inner nuclear layer (INL), which hosts horizontal, bipolar and amacrine cells and is located in between the RGL and ONL. In this way, we wanted to assess how nuclear packing changes during the transition from the retinal PSE to the laminated retina.

To quantitatively track the changing nuclear morphologies of emerging neurons, we trained a different StarDist-3D model on a manually annotated dataset that specifically included annotations from the nuclear layers of laminated retinas between 48 hpf, by which time neuronal lamination has started, and 80 hpf, when lamination is complete (Fig. 4 A, B; Fig. S1 E; Table 4). Across all three layers, nuclear volume progressively decreased with time (Fig. 4 G). Nuclear aspect ratios, however, were differently distributed (Fig. 4 C, D), especially across the distributions of the shorter axis lengths (Fig. S4 C, I). The distribution of the major axis lengths did not significantly vary across the three layers and progressively decreased over developmental time (Fig. S4 C, H). As a result, nuclei in the RGL and INL became more spherical, while photoreceptor precursors in the ONL featured an elongated rod-like shape (Fig. 4 C,D). As nuclei assumed different shapes and sizes depending on their neuronal fate, the linear correlation between number of contacts and nuclear aspect ratios was lost across all three layers compared to nuclei in the PSE (Fig. S5 G). This, together with the fact that the mean internuclear distance did not significantly change during neuronal lamination (Fig. S5 J), suggested that nuclei had reached their closest packing configuration by 80 hpf.

**Fig. 4.**
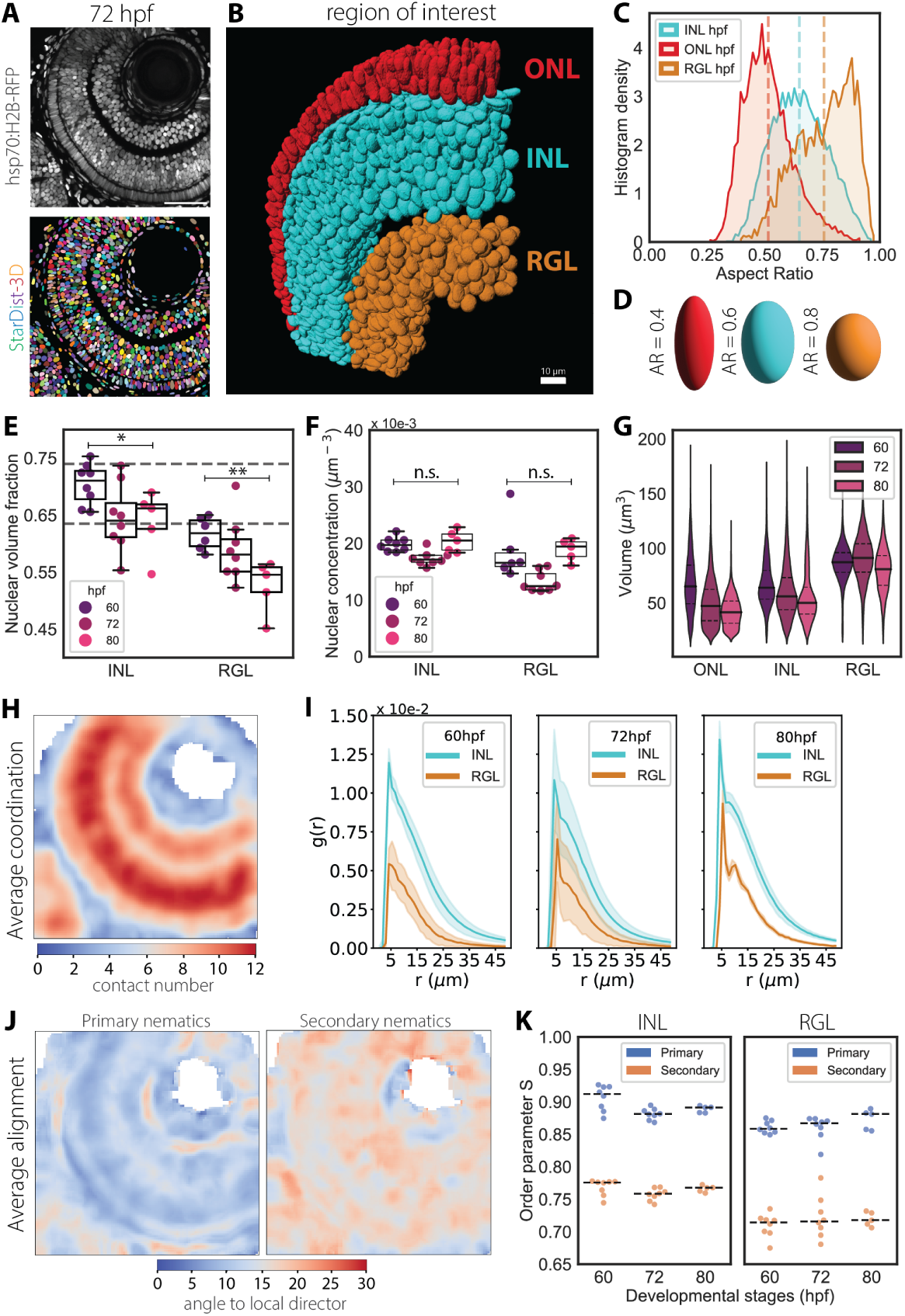
Nuclear packing reaches theoretical limiting density as nuclear shape and sizes are remodelled during neuronal lamination. **A)** Representative images of the laminated retina at 72 hpf (upper) and corresponding instance segmentation using StarDist-3D model (bottom). Nuclei are labelled with Tg(hsp70:H2BRFP). Scale bar is 50 *µ*m. **B)** 3D-rendering of 72 hpf ROI from panel (A), color-coded for distinct nuclear layers. Scale bar is 10 *µ*m. **C)** Histogram density distribution for nuclear aspect ratios at 72 hpf. Segmented lines indicate the mean values for each distribution. **D)** Schematics of prolate ellipsoids having aspect ratios as shown in panel (C). **E)** Nuclear volume fractions of nuclei in the inner nuclear layer (INL) and retinal ganglion cell layer (RGL). 60 hpf, N = 7; 72 hpf, N = 8; 80 hpf, N = 5 embryos. **F)** Concentration of nuclei found within the ROIs of panel (E). P values for alpha = 0.01: 0.01 *<* * *<* 0.05; ** *<* 0.01; n.s. not significant; from unpaired two-tailed Student’s t test. **G)** Nuclear volume distributions. See table for test statistics.H) Median intensity projections of image in panel (A) reporting of the average coordination number for nuclei within a window of size 10×10×10 *µ*m. **I)** Radial distribution function g(r) in INL and RGL over development. Solid lines correspond to mean values, shaded area indicate the standard deviation from the mean. **J)** Median intensity projections of image in panel (A) reporting of the local angle of misalignment for the primary (left) and secondary (right) nematic axes within a window of size 10×10×10 *µ*m. **K)** Order parameter S.

To explore how the local packing environment changed over lamination, we performed the same analysis of nuclear packing order as done for the proliferative phase. Because photoreceptors in the ONL are organized in a contiguous monolayer at the tissue boundaries of the retina and are visibly arranged in a regular linear pattern (Fig. 4 A, B), we focused our analysis on the INL and the RGL. Visual inspection of smoothed images confirmed that nuclei in the delaminating retina reached higher contact numbers compared to the RNE at 48 hpf, with the INL featuring the closest configuration (Fig. 4 H). In the INL, nuclei reached their highest packing densities around 60 hpf (Fig. 4 E), at values approximating the *ϕ_e_* ≈ 0.74 value, known to be the RCP density for fully aspherical ellipsoids [45]. Between 60 hpf and 72 hpf, both nuclear packing and nuclear concentration decreased in the INL and RGL, possibly as a consequence of tissue growth and positional ordering (Fig. 4 E, F). Correlations between volume fractions and nuclear concentration differed when the INL or RGL were compared to the retinal PSE (Fig. S4 B, D), suggesting a distinct proliferation and differentiation state of cells in these layers, where retinal ganglion cells mature earlier than other neurons [53]. Furthermore, we calculated the RDF of nuclei in the INL and RGL to determine whether nuclear arrangements were shifting to more regular structures during neuronal lamination. Strikingly, we found discrete peaks in the RDF by 80 hpf both in the INL and in the RGL, indicating the highest probability to find a nucleus within 5 and 10 *µ*m radius (Fig. 4 I). This suggests that ordered, periodic arrangements of nuclei emerge during neuronal lamination. Moreover, when we analysed the alignment of nuclei across the three neuronal layers, we found that nuclei maintained a strong orientation along their primary nematic axis even at later stages of development (Fig. 4 J, K).

We conclude that differentiation of neuronal precursors coincides with changes in nuclear shape, alongside the formation of a more ordered, crystal-like arrangement in the INL and RGL. In the ONL, each photoreceptor is positioned adjacent to its neighbour to form a densely packed monolayer of elongated nuclei arranged in a regular pattern [54]. Despite changes in morphology and position, nuclei of neurons remain aligned along the radial axis of the neural retina, similar to the alignment reported in the proliferative stage. This indicates that orientational order is established early on in development and is conserved in the laminated retina during the emergence of crystalline positional order.

**Table 2.** Nuclear volume, axes lengths and aspect ratios of nuclei in the laminated retina between 60 and 80 hpf. Mean and standard deviation are shown for each stage and nuclear layer. Mann-Whitney tests between previous and following stages were performed on nuclear volume distributions and aspect ratio distributions across nuclear layers, e.g. comparing distributions in the INL between 60 hpf and 72 hpf and between 72 hpf and 80 hpf. P-values are shown for each comparison.

| | | Nuclear volume($\mu\text{m}^3$) | | |
| --- | --- | --- | --- | --- |
| Layer | Stage | $\mu \pm \sigma$ | H0: same distribution | p value |
| ONL | 60 hpf | 56.77 ± 29.12 | – | – |
| | 72 hpf | 45.89 ± 22.80 | (60 hpf) = (72 hpf) | p ≪ 0.0001 |
| | 80 hpf | 36.80 ± 16.56 | (72 hpf) = (80 hpf) | p < 0.001 |
| INL | 60 hpf | 59.11 ± 23.33 | – | – |
| | 72 hpf | 58.39 ± 24.37 | (60 hpf) = (72 hpf) | p ≪ 0.0001 |
| | 80 hpf | 53.46 ± 24.45 | (72 hpf) = (80 hpf) | p < 0.001 |
| RGL | 60 hpf | 79.41 ± 19.44 | – | – |
| | 72 hpf | 90.11 ± 21.24 | (60 hpf) = (72 hpf) | 0.013 |
| | 80 hpf | 79.58 ± 20.19 | (72 hpf) = (80 hpf) | p ≪ 0.0001 |
| | | Nuclear axis lengths ($\mu\text{m}$) | | |
| Layer | Stage | A ($\mu \pm \sigma$) | B ($\mu \pm \sigma$) | C ($\mu \pm \sigma$) |
| ONL | 60 hpf | 7.11 ± 1.60 | 4.28 ± 0.99 | 3.41 ± 0.86 |
| | 72 hpf | 7.02 ± 1.53 | 3.94 ± 0.80 | 3.12 ± 0.66 |
| | 80 hpf | 6.80 ± 1.39 | 3.54 ± 0.62 | 2.89 ± 0.55 |
| INL | 60 hpf | 6.89 ± 1.29 | 4.44 ± 0.80 | 3.63 ± 0.70 |
| | 72 hpf | 6.47 ± 1.09 | 4.55 ± 0.76 | 3.73 ± 0.73 |
| | 80 hpf | 5.98 ± 1.05 | 4.44 ± 0.73 | 3.70 ± 0.72 |
| RGL | 60 hpf | 6.82 ± 1.02 | 5.15 ± 0.59 | 4.32 ± 0.59 |
| | 72 hpf | 6.80 ± 1.00 | 5.44 ± 0.54 | 4.70 ± 0.59 |
| | 80 hpf | 6.38 ± 0.87 | 5.18 ± 0.51 | 4.57 ± 0.57 |
|  |  | Aspect ratio (A/(B+C)) |  |  |
| Layer | Stage | ($\mu \pm \sigma$) | H0: same distribution | p value |
| ONL | 60 hpf | 0.55 ± 0.13 | – | – |
| | 72 hpf | 0.50 ± 0.10 | (60 hpf) = (72 hpf) | p < 0.001 |
| | 80 hpf | 0.48 ± 0.09 | (72 hpf) = (80 hpf) | p < 0.001 |
| INL | 60 hpf | 0.59 ± 0.11 | – | – |
| | 72 hpf | 0.65 ± 0.11 | (60 hpf) = (72 hpf) | p ≪ 0.0001 |
| | 80 hpf | 0.69 ± 0.11 | (72 hpf) = (80 hpf) | p < 0.001 |
| RGL | 60 hpf | 0.70 ± 0.11 | – | – |
| | 72 hpf | 0.75 ± 0.10 | (60 hpf) = (72 hpf) | 0.011 |
| | 80 hpf | 0.77 ± 0.10 | (72 hpf) = (80 hpf) | 0.1 |

### Modelled material properties of the developing RNE switch between a cellular-dominated and nuclear-dominated states, affecting tissue shape

Since neuronal lamination correlates with repositioning and shape remodelling of nuclei from nematically ordered to a more crystal-like arrangement, we asked whether neuronal lamination itself could allow for more nuclei and cells to be packed within the tissue without resulting in a premature rigidity transition. To theoretically explore the effects that the nuclear packing environment can have on the mechanical state of the developing RNE, we took inspiration from a previously published continuum formulation of an epithelium growing under confinement [55]. We followed this approach while also incorporating internal states corresponding to cellular-dominated and nuclear-dominated mechanical regimes (Fig. 5 A). In this way, we could evaluate the separate potential contributions to the compression and bending moduli of the two-state modelled growing epithelium (see Theory Supplement). Based on this model, we observe a sudden increase in the modelled growing epithelium’s instability to buckling as it approaches the jammed, nuclear-mechanics dominated internal state which corresponds to a sudden decrease in the critical buckling strain (Fig. 5 B). Therefore, according to this model, regulation of the internal state of the RNE directly affects tissue growth and shape and sets a mechanical constraint on the proliferation of the hemispherical RNE. In this context, we speculate that the onset of neuronal lamination could prevent the retinal PSE from reaching the nuclear jamming transition prematurely, when single cells and nuclei are actively moving through the tissue to reach their final positions [36]. In this way, neuronal lamination would promote the transition from positional disorder to order, thereby keeping the RNE in a cellular-dominated internal state and preventing the onset of a buckling instability. This, in turn, would enable the correct growth and scaling of the optic cup.

**Fig. 5.**
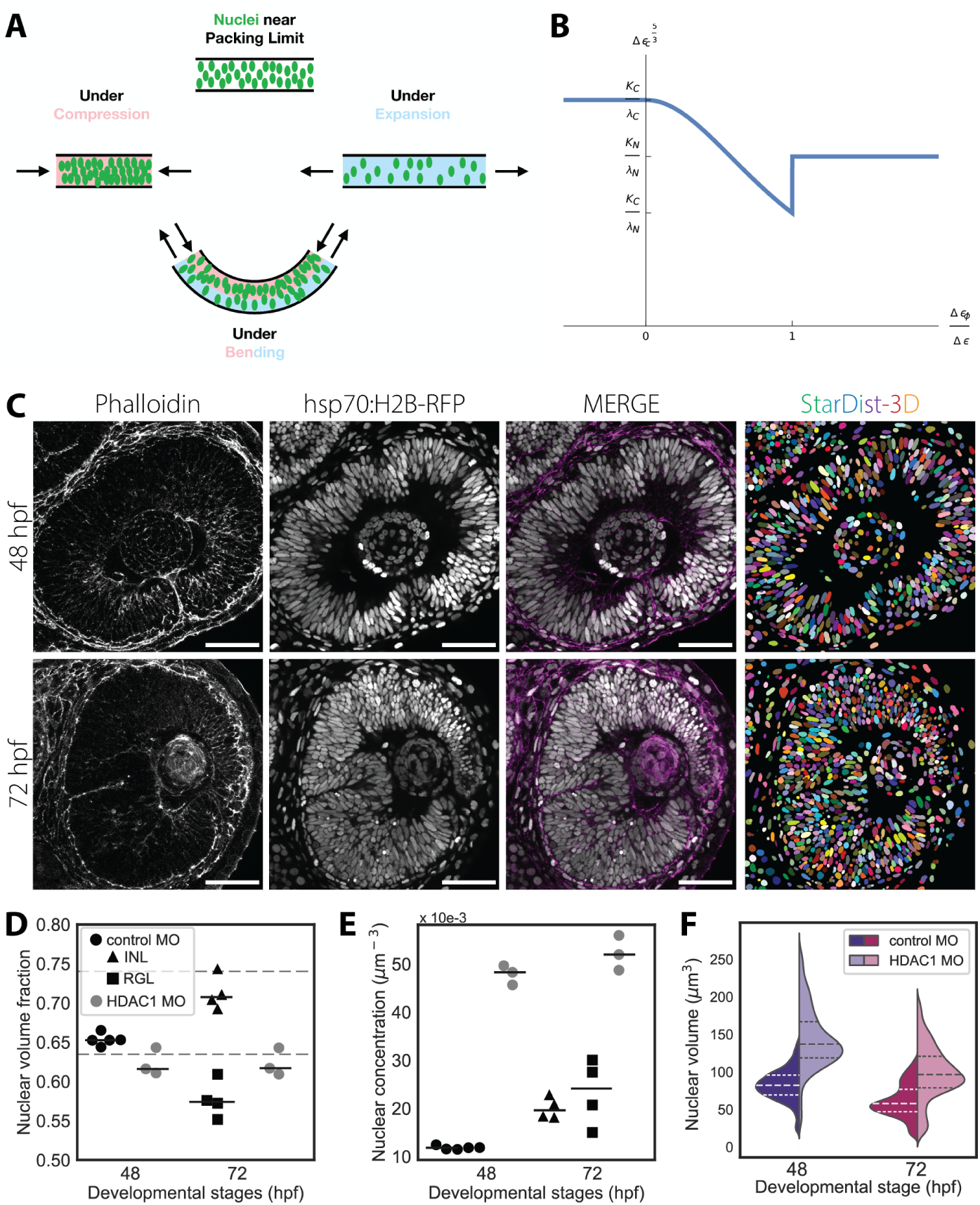
Nuclear-dominated mechanical state leads to buckling when delamination does not occur. **A)** Diagram of the interplay of the internal packing state of the tissue’s nuclei with externally imposed deformations: compression, expansion, and bending. Compression can drive the mechanics from cellularly dominated to nuclearly dominated while expansion is the reverse. Bending creates a mix of both. **B)** The critical strain as a function of Δ*∈_ϕ_*/Δ*∈*, where Δ*∈* is the excess strain and Δ*∈_ϕ_* is the excess strain to the jamming transition boundary. Note that *λ_c_* and *λ_n_* are the cellular and nuclear compression stiffness, *K_c_* and *K_n_* are the cellular and nuclear-associated bending rigidities. C) Representative confocal sections of 48 hpf (top row) and 72 hpf (bottom row) retinas in Hdac1 morphants with corresponding StarDist-3D segmentation predictions. Nuclei are marked with Tg(hsp70:H2B-RFP), cell membranes with phalloidin. Scale bar is 50 *µ*m. D) Volume fractions occupied by nuclei in ROIs. Control morphants (MO), N = 5 embryos; HDAC1 MO, N = 3 embryos. Comparison at 48 hpf, n.s.; comparison at 72 hpf (INL), p value *<* 0.01; comparison at 72 hpf (RGL), n.s. for unpaired Student’s t-test. E) Concentration of nuclei found within the ROIs of panel (D). comparison at 48 hpf, p value *<* 0.00001; comparison at 72 hpf (INL), p value *<* 0.00001; comparison at 72 hpf (RGL), p value *<* 0.01 for unpaired Student’s t-test. F) Nuclear volume distributions. See table for test statistics.

### Blocking neurogenesis interferes with nuclear shape and size changes and leads to buckling of the retinal PSE

Based on our one-dimensional model of epithelia growing under confinement, we hypothesized that timely control of neuronal lamination ensures that the developing RNE does not approach the nuclear jamming transition, which would subsequently lead to deformations of the tissue. In this scenario, timely progression towards a laminated state could shift the threshold to entry into the nuclear-dominated mechanical state to higher nuclear packing densities and allow correct growth and shape scaling. To test this, we probed the limits of nuclear packing in the RNE by perturbing neuronal lamination and thereby keeping the retina in a PSE-like arrangement for longer.

To impair neuronal lamination, we blocked neurogenesis and promoted sustained proliferation in the RNE, using a previously established Histone Deacetylase 1 (HDAC1) morpholino knock-down (KD) approach [21]. HDAC1 suppresses Wnt and Notch signalling in the RNE, therefore regulating the balance between proliferation and neuronal differentiation of multipotent progenitors [56, 57]. When HDAC1 is impaired, progenitor cells do not exit the cell cycle, continue dividing and do not differentiate into neurons, thereby blocking neuronal lamination [21, 56, 57].

Using our previously established image analysis pipeline, we observed that progenitor cells in HDAC1 morphants are still organized into a PSE-like structure even at stages when the retina is already laminated in the control scenario at 72 hpf. The overall shape of the RNE starts to be visibly perturbed after 48-50 hpf, with the retinal PSE undergoing buckling [21, 56, 57]. We used the tg(lama1:lama-GFP) transgenic line that labels the surrounding basal lamina to segment the optic cup volume and determine whether it differed between morphants and controls (Fig. S6 A,B). Despite continued proliferation, our segmentation showed that the volume of the optic cup was smaller in morphants than in controls (Fig. S6 E). Thus, eyes in HDAC1 morphants failed to reach a normal organ size at stages when the retina is normally organized in a layered architecture.

Given that the morphant optic cup volumes were reduced and multipotent progenitors could not exit the cell cycle to differentiate into postmitotic neurons, we probed nuclear packing densities in the RNE upon HDAC1 KD at 48 and 72 hpf, i.e. before and after buckling. Predictions obtained with our StarDist-3D model had to be manually curated and were analysed using our image analysis pipeline (Fig. 5 C). Nuclei in HDAC1 KD RNE were bigger and extremely elongated along their major axis prior to buckling (Fig. 5 F; Fig. S6 C, F, G; Table 3). The extreme elongation of these nuclei suggested that they may have been deformed by increasing lateral compressive stresses building up prior to the onset of any buckling event. Nuclear size was reduced only upon buckling of the PSE at 72 hpf (Fig. 5 F) when nuclei increased their aspect ratios by shortening of the major axis lengths, but nevertheless not reaching control ratios (Fig. S6 F, G; Table 3). These results indicate that retinal progenitor cells in HDAC1 morphants could not fully adjust their nuclear size and shape as seen for progenitor cells in control embryos upon lamination onset. Despite their extremely elongated shapes, nuclei in the HDAC1 morphants did not contact more neighbours than in the laminated retina of control embryos (Fig. S6 H), ssuggesting that the nuclei had reached the closest packing configuration for this PSE-like arrangement. Measurements of the nuclear volume fraction in HDAC1 morphants did not increase between 48 and 72 hpf and did not reach *ϕ_s_* as in controls, even though nuclear concentrations were higher in HDAC1 morphants (Fig. 5 D, E). This suggests that, in controls, the retinal PSE is approaching its closest packing density at around 48 hpf, when the RNE is normally transitioning to a layered structure, and is approaching a nuclear jamming threshold that could lead to buckling of the tissue, as suggested by our toy model (Fig. 5 A, B).). In HDAC1 morphants, the retinal PSE deformed and buckled between 48 hpf and 72 hpf, while the retinal pigmented epithelium (RPE), which covers the apical surface of the RNE, did not buckle with it (Fig. S6 D). Nuclei in the RPE showed elongated morphology where the RNE was not bending and, conversely, featured rounder morphologies above folded portions of the retina. Therefore, only the retinal PSE was undergoing shape deformations in the optic cup of HDAC1 morphants.

**Table 3.**
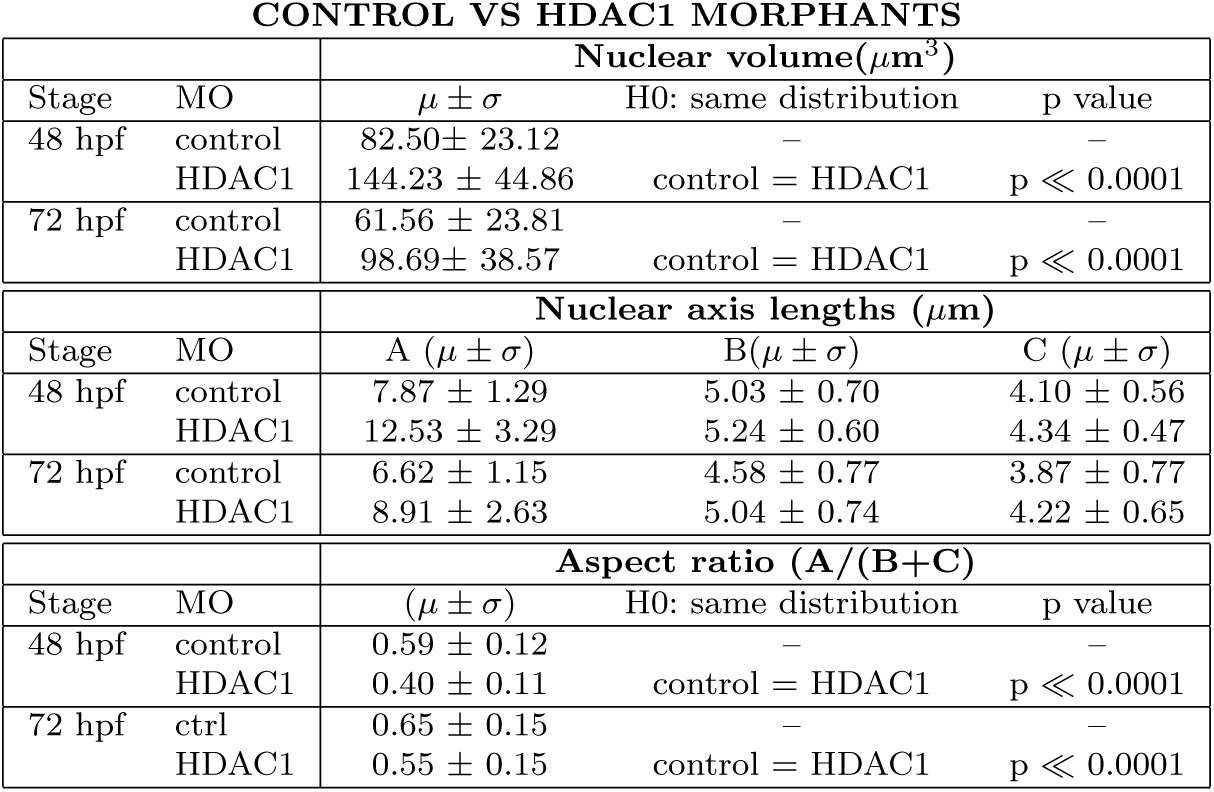
Nuclear volume, axes lengths and aspect ratios of nuclei in the retina of embryos treated with either control or HDAC1 morpholinos (MO). Mean and standard deviation are shown for each stage and nuclear layer. Mann-Whitney tests between previous and following stages were performed on nuclear volume distributions and aspect ratio distributions between control and treated samples.

Furthermore, the phenotype of HDAC1 morphants closely resembles the buckling dynamics predicted by our theoretical model. This suggests that, upon disruption of neurogenesis onset in HDAC1 morphants, the RNE transitions to a nuclear-dominated mechanical state, where the critical buckling strain decreases, thus facilitating tissue buckling. Our findings further indicate that tissue buckling releases some of the compressive stresses that affected nuclei in the RNE, while other tissues, like the RPE, were not as affected. In contrast, it was shown that loss of HDAC1 activity in the hindbrain does not lead to tissue buckling [58], supporting our hypothesis that buckling results from deregulation of nuclear packing configurations in densely crowded PSE, like the RNE. Overall, these results from theory and experiment suggest that the remodeling of nuclear shape and size during neuronal lamination are linked to the mechanical state of the tissue, therefore affecting growth and shape of the retina.

## Discussion

In this study, we characterized nuclear packing in vivo and over development in the retinal and hindbrain neuroepithelia using deep-learning methods to achieve accurate 3D segmentation of nuclei. In this way, we generated new insights into how nuclear packing progressively transitions to ordered nuclear arrangements and how this can influence morphogenesis. We find that in both tissues nuclear packing increases but to different degrees, with the nuclear packing in the retina approaching limiting packing fractions. Nevertheless, in both retinal and hindbrain PSE nuclei are arranged in a nematic-like order indicating that this is the conserved tissue configuration set by pseudostratification. The strong alignment of nuclei along the apicobasal cell axis throughout development could facilitate the radial nuclear translocations that characterize proliferating PSE. Moreover, inclusion of a nematic-like order during ongoing nuclear packing increase suggests that both neuroepithelia are possibly transitioning to a glass-like phase, with subsequent jamming transition of the tissue, as recently suggested [33]. This nematic-like order could represent one of the advantages that characterize pseudostratification.

We show that in the retina, nuclear packing progressively transitions to a crystalline order upon neuronal lamination due to remodelling of nuclear sizes and shapes. Interestingly, the orientational order of nuclei along the tissue radial axis, which is established by pseudostratification, prevails at these stages. We also find that timely reorganization of nuclear packing from a pseudostratified to a layered arrangement is important for maintaining the shape of the optic cup and enables tissue growth while sustaining hemispherical morphology.

To obtain these results, we performed, to our knowledge, the first 3D instance segmentation of nuclei in proliferating PSE over development. This enabled us to accurately detect nuclei within volumetric imaging datasets of crowded tissues. Nuclei of neuroepithelial cells were found to be positionally disordered within the retinal PSE, and featured long-range alignment along the apicobasal axis, i.e. orientational order. This organization resembles the organization of anisotropic particles in a nematic liquid crystal and enables closer configurations of nuclei, while the RNE remains positionally disordered. The particular biaxial nematic ordering of nuclei found in the RNE is conserved also in another PSE, the developing zebrafish hindbrain, which features a tubular tissue shape and different regulation of growth and patterning. Despite differences in proliferation, neurogenesis onset and, importantly, packing density, both neuroepithelia follow the same nuclear ordering rules during their early morphogenesis. This suggests that the conservation of pseudostratification could be linked to the properties of this biaxial nematic-like organization.

A nuclear jamming transition has been recently proposed in 2D simulations of epithelia during development [33]. Here, neuroepithelial cells were shown to become progressively constrained when the nuclear volume fraction increased and nuclear anisotropy decreased. In that study, comparison of these *in silico* findings to 2D optical slices of the zebrafish retina suggested that nuclei in the INL reached a jamming transition boundary between 55 and 72 hpf, similarly to what is observed here. Additional to the findings in this 2D analysis, our 3D analysis of nuclear packing showed how nuclear arrangements in the RNE transition from nematic order to a fully crystal-like periodic order, while still retaining long-range alignment. We showed that the observed decrease in both nuclear size and shape anisotropy is downstream of neuronal differentiation and that timely onset of neuronal lamination enables correct tissue growth and morphology. At the same time, neuronal lamination avoids the onset of a compressive buckling instability that is strongly enhanced by the approach to a nuclear-dominated mechanical state. Taken together, this demonstrates how nuclear shape anisotropy and packing can drive tissue organization and support the existence of a nuclear jamming transition in developing neuroepithelia. These findings should be further compared to other studies in which the role of nuclear shape and positioning in remodelling tissue topologies of other developing epithelia, like the germband of *Drosophila* embryos [59], was recently explored.

The presented 3D analysis allowed us to characterize nuclear shape and size changes over the course of retinal development and showed that the timing of the onset of neuronal lamination coincided with Random Close Packing densities. When we probed the limits of the retinal PSE upon perturbation of neurogenesis, we found that nuclear packing in the RNE could not reach higher volume fractions without transitioning to a layered tissue. This suggests that in the RNE the limiting nuclear packing for a PSE is reached around the time of neuronal lamination, at least in the vertebrate retina. When blocking neurogenesis and neuronal lamination by the HDAC1 KD, severe tissue deformations were observed. This is in line with several theoretical and experimental studies proposing that proliferation of epithelia under confinement can induce buckling instabilities, i.e. without the requirement of active bending mechanisms [55, 60, 61]. Epithelial cells can partially accommodate compressive stresses by increasing their height until the critical strain for the compressive instability is reached, after which the tissue buckles [60]. Progenitor cells in HDAC1 morphants are characterized by extremely elongated nuclei prior to buckling of the RNE, whilst they fail to increase their cell height [21]. Meanwhile, the RPE is stretched over the RNE, with the exception of those portions where the retina has buckled. We speculate that the RPE could confine the growth of the retina and therefore contribute to the buckling phenotype from the outside. In conclusion, we propose that the strong nuclear anisotropy we observe upon HDAC1 KD is due to the compressive stresses generated by the proliferating cells under confinement, until these stresses cannot be accommodated any longer for dense nuclear packing in a PSE-like arrangement, leading to tissue buckling. This is in line with the predictions from our one-dimensional continuum model that shows that the approach to a nuclear-mechanics dominated state further facilitates tissue buckling. Based on this, we propose that neuronal lamination avoids the buckling instability in the retina by keeping the tissue sufficiently in the looser, cellular-mechanics dominated state.

Over the course of neuronal lamination, nuclei of neuronal precursors change their shape, size and position, leading to the emergence of crystal-like arrangements. Interestingly, we find that most nuclei across the three nuclear layers are still aligned along the radial axis of the tissue, which coincides with the physiological direction of light propagation. Because photons need to pass through the whole thickness of the retina to reach the photoreceptors, correct and precise vision in vertebrates is aided by minimal light scattering. It has been proposed that scattering is reduced by the light-guiding capabilities of the Müller Glia [62, 63] and by precise spatial organization of heterochromatin of rod photoreceptors in nocturnal animals [64, 65]. Our study indicates that nuclei in the retina are arranged with orientational and crystalline order, which could further enhance tissue transparency and light propagation, as previously suggested by simulations of ordered nuclear arrangements [64, 65].

In conclusion, our study shows that pseudostratification establishes a nematic-like order early on during organogenesis and that the emergence of crystal-like nuclear arrangements enables close nuclear packing of newly formed neurons without affecting tissue shape. These insights were only possible thanks to 3D nuclear segmentation over the entire proliferative and neurogenic phases. Since zebrafish is characterized by a particularly fast-paced embryogenesis, it will be interesting to compare these findings in the zebrafish RNE to other teleosts and vertebrates that develop over longer timescales. For instance, future studies could dissect the distribution of forces and stresses within proliferating and crowded neuroepithelia and compare the timely regulation of proliferation and neurogenesis across various vertebrate species. This will inform us about the mechanisms that control nuclear packing order and size and shape of the retina over different developmental time windows.

## Methods

### Zebrafish husbandry

The experimental work reported in this article was performed on *Danio rerio*, i.e. zebrafish, embryos aged between 24 and 120 hours post fertilization (hpf). Wild-type (AB and Tupfel long-fin (TL) strains) and transgenic fish were bred and maintained in a recirculation life support system (Tecniplast) with the following parameters: 28 °C, pH 7.0, conductivity 1000 *µ*S/cm, 14L:10D light:dark cycle. Up to juvenile stage, fish were fed with a combination of saltwater rotifers (Brachionus plicatilis) and processed diet (Gemma 150, Skretting). Adult fish were fed with a combination of live food (Artemia salina) and commercial processed dry food (Gemma 300, Skretting).Embryos used for experimental work were raised at 21°C, 28.5°C, or 32°C in E3 medium supplemented with 0.2 mM 1-phenyl-2-thiourea (10107703, Acros Organics) from 8 hpf to prevent pigmentation. Embryos were staged according to. Kimmel et al. 1995 [66]. Anesthesia was performed by supplementing the E3 medium with 0.04% tricaine methanesulfonate (MS-222, 1004671, Pharmaq) before live imaging. All animal work was conducted in accordance with institutional standard operating procedures under the licensing of the DGAV (Direcção Geral de Alimentação e Veterinária, Portugal) and in accordance with the European Union directive 2010/63/EU and with the Portuguese Decree Law n° 113/2013.

### Transgenic lines

To visualize all nuclei in the retina, the Tg(*β*-act:H2A-GFP)[67] and the Tg(hsp70:H2B-RFP) [68] line were used. Tg(atoh7:gap-GFP) and Tg(atoh7:gap-RFP) zebrafish transgenic lines were used to identify Atoh7+ progenitors and Atoh7+ neurons [69]. The Tg(lama1:lama1-sfGFP) was used to visualize the outline of the optic cup [70].

### Heat shock of embryos

Transgenic lines expressing constructs under the heat shock promoter (hsp70) were incubated in water bath set to 37.5°C for at least 20 min to drive the expression of the construct. In the case of Tg(hsp70::H2B-RFP), embryos were heat-shocked for 20-30 minutes and imaging was started around 3-4 hours after. For embryos analysed at stages beyond 36 hpf, the heat shock was performed twice to enhance the signal: one time the day before the experiment and a second time 4 hours before imaging or fixing. For all the mentioned transgenic lines, controls were defined as embryos from the transgenic line that were heat-shocked as described here, but did not express the construct of interest.

### Morpholino experiments

All morpholinos were purchased from Gene Tools, LLC. Morpholino targeting HDAC1 (5’-TTGTTCCTTGAGAACTCAGCGCCAT-3’) was injected at 0.5 ng per embryo, together with the morpholino targeting p53 (5-GCGCCATTGCTTTGCAAGAATTG-3), which was added to the injection mix (0.75 ng/ embryo) to alleviate cell death possibly resulting from the off-target effects of the morpholino. Control embryos were injected with a scrambled morpholino (5’-CCTCTTACCTCAGTTACAATTTATA-3’) injected at 0.5 np per embryo, together with the p53 morpholino.

### Whole mount immunofluorescence

All immunostainings were performed on whole-mount embryos fixed overnight in 4% paraformaldehyde (043368-9M, Thermo Fisher Scientific) in PBS 1X at 4°C as previously described [47, 53]. Embryos were washed from 3 to 5 times for 10 min in PBS-Triton 0.8% (28817295, VWR) and permeabilized in 0.25% Trypsin-EDTA (sc391060, Santa Cruz Biotechnology) on ice for different time periods depending on the developmental stage: 10 min for 24 hpf, 28 hpf and 36 hpf; 12 min for 42 hpf; 15 min for 48 hpf, 60 hpf, 72 hpf and 120 hpf. Embryos were then washed for 30 min on ice in PBS-Tr 0.8% on shaker and then incubated in blocking solution, i.e. 10% goat serum (16210064, Gibco) in PBS-Triton 0.8%, for 3 hours at room temperature. Embryos were then incubated with the following primary antibodies for 72 hours at 4°C on shaker: anti-GFP, 1:100 (Proteintech, 50430-2-AP); histone H3 phosphorylated on S28, 1:500 (Abcam, ab10543); anti-HuC/HuD, 1:250 (Invitrogen, A-21271). Embryos were washed 3 times for 30 min with PBS-Triton 0.8% and then incubated for 24-48 hours with fluorescently labelled secondary antibodies (Molecular Probes). In the case of the Tg(hsp70::H2B-RFP) transgenic line, embryos were incubated in antibody solution with RFP booster (1:200, ChromoTek, rba594) to enhance the fluorescence signal of the expressed RFP. Phalloidin conjugated with Alexa Fluorophore 405 (1:100, Biotium; 00034-T), Alexa Fluorophore 488 (1:50, Life Technologies; A12379) and Alexa Fluorophore 647 (1:50, Cell Signalling; 8940) were used to label F-Actin in all cells. Finally, embryos were washed several times with PBS-Triton 0.8% in ice on shaker and stored in PBS at 4°C until imaging.

### Microscopy

#### Laser scanning confocal microscopy

Dechorionated, fixed and immunostained samples were imaged with a Zeiss LSM980 Airyscan2 inverted microscope, equipped with two PMT and one GaAsP detector, using a 40X/1.1 C-Apochromat water immersion objective from Zeiss. Embryos were mounted in 0.70% low-melting agarose in glass-bottomed dishes (35 mm, MatTek Corporation) and imaged at room temperature. The dataset used for nuclear segmentation was acquired with Airyscan CO-8Y mode and processed using the Airyscan processing methods for 3D datasets. The microscope was operated using the proprietary ZEN Blue v3.3 software.

#### Light sheet fluorescence microscopy

Dechorionated live tg(lama1:lama1-sfGFP) embryos were mounted in 1 mm glass capillaries in 0.6% low-melting agarose. The sample chamber was filled with E3 medium containing 0.01% MS-222 (Sigma) and 0.2 mM PTU (10107703, Acros Organics). Imaging was performed on a Zeiss Lightsheet Z.1 microscope equipped with two PCO Edge 4.2 sCMOS cameras (max 30 fps with 2048×2048 pixels) and with a 20x/1.2 Zeiss Plan-Apochromat water-immersion objective. Z-stacks spanning the entire retinal neuroepithelium were acquired with 1 *µ*m optical sectioning with double-sided illumination mode. The microscope was operating using the proprietary ZEN Black v3.0 software.

### Image analysis

#### Segmentation of the optic cup

3D segmentation of the basement membrane was performed using the FIJI [71] plugin for 3D segmentation LimeSeg (v 0.4.2) [72]. The outline of the optic cup was marked by the tg(lama1:lama1-sfGFP) line. Several regions of interest (ROIs) where drawn inside the volume of the optic cup across the z-stack. End-points were defined by a single point. All ROIs were saved in the FIJI ROI manager and sorted according to their position along the z axis. The ‘Skeleton Seg’ approach in LimeSeg was used with the following parameters: D0 (minimal diameter of smallest object in pixel) equal to 16; F pressure (default pressure) of 0.0099; Range in d0 units equal to 1; number of integration steps of −1. The resulting 3D mesh of the object’s surface was saved in ply format and a python script was coded to convert this mesh into a labelled mask in tiff format (the script can be found in the github repository of this study). The resulting labelled objects were then manually corrected using the label tools in Napari (v0.4.18) [73].

#### 3D instance segmentation of nuclei

Deep-Learning methods generally requires vast amounts of pixel-wise annotated ground truth data for training. For this reason, we manually annotated up to 16 ground-truth 3D cropped images of various sizes, at least 50×260×230 pixels having voxels size equal to zyx = [0.24, 0.102, 0.102] in *µ*m, taken from several datasets representing various developmental stages. The stacks sizes were defined by the volume of the nuclei, since each crop needed to be big enough to contain fully visible nuclei, i.e. nuclei not touching the image border. Manual annotation was performed using the label tools in Napari.

Using the aforementioned annotated stacks, we trained two StarDist-3D models: one model (model A) was trained on 16 ground-truth stacks representing stages between 24 and 72 hpf, while the other model (model B) was trained on 14 stacks representing later stages of the laminated retina, i.e. between 48 and 80 hpf. Each stack contained between 10 and up to 110 entire labelled objects, i.e. nuclei that were not touching the image border. In both training datasets, the z step was twice the voxel size in xy and this anisotropic factor was kept in the imaging dataset. The voxel size used was comparable to the voxels size in the training datasets. To artificially increase the size of the training dataset and make the model more robust to pixel intensity fluctuations, the training dataset was augmented by adding synthetical noise on all pixels and rotating, flipping, transforming each labelled image. The following parameters were used to optimize the neural network for the training: number of rays, 256; grid size, [1, 4, 4]; anisotropy, [4.17, 1.27, 1.0]; backbone, u-net; u-net pool, [2, 4, 4]; train patch size, [32, 128, 128]; epoch; 400; steps per epoch, 100; train learning rate, 0.003. The script coded to perform the StarDist-3D training on our data can be found in the github repository of this study and can be reused under open-source licence conditions.

In order to evaluate the trained models, predictions were run on another manually annotated dataset that had never been shown to StarDist-3D during training. Table 4 shows the accuracy, precision and recall for several Intersection over Union (IoU) thresholds *τ* for test datasets representing different developmental stages, i.e. 30, 48 and 72 hours post fertilization (hpf). The number of test ground-truth crops per stage was, respectively : N = 2; N = 3; N = 4. The total number of nuclei per stage, respectively: n = 272; n = 225; n = 647. StarDist-3D model A was tested on datasets from 30 and 48 hpf embryos, while model B was tested on dataset from 72 hpf embryos. Since StarDist-3D is well suited for objects that can be represented by star-convex polyhedra, such as in the case of round and elliptical nuclei, our models could not produce accurate predictions of extremely elongated nuclei. This was the case for some of the elongated nuclei in wild-type retinas at 24 hpf and in HDAC1 morphants. Therefore, the imaging dataset of retinas from these samples was segmented using our StarDist-3D model and the predictions were manually curated using Napari (v0.4.18) and the annotation tool from Segment Anything for Microscopy (SAM, v0.3.0) [74].

**Table 4.**
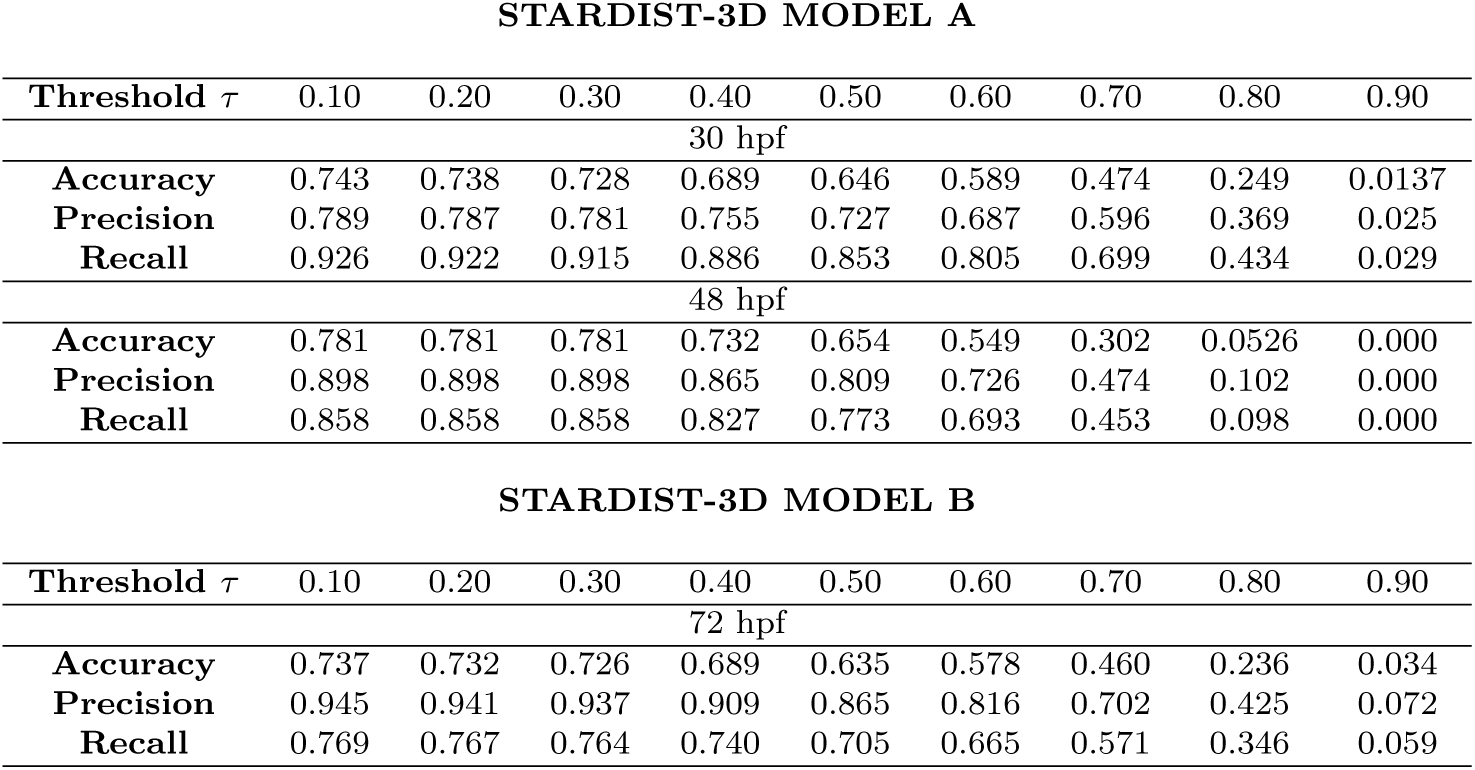
Evaluation of StarDist-3D models using crops of imaged nuclei from embryos staged at 30, 48 and 72 hpf.

**STARDIST-3D MODEL A**
| Threshold $\tau$ | 0.10 | 0.20 | 0.30 | 0.40 | 0.50 | 0.60 | 0.70 | 0.80 | 0.90 |
| --- | --- | --- | --- | --- | --- | --- | --- | --- | --- |
| 30 hpf |  |  |  |  |  |  |  |  |  |
| **Accuracy** | 0.743 | 0.738 | 0.728 | 0.689 | 0.646 | 0.589 | 0.474 | 0.249 | 0.0137 |
| **Precision** | 0.789 | 0.787 | 0.781 | 0.755 | 0.727 | 0.687 | 0.596 | 0.369 | 0.025 |
| **Recall** | 0.926 | 0.922 | 0.915 | 0.886 | 0.853 | 0.805 | 0.699 | 0.434 | 0.029 |
| 48 hpf |  |  |  |  |  |  |  |  |  |
| **Accuracy** | 0.781 | 0.781 | 0.781 | 0.732 | 0.654 | 0.549 | 0.302 | 0.0526 | 0.000 |
| **Precision** | 0.898 | 0.898 | 0.898 | 0.865 | 0.809 | 0.726 | 0.474 | 0.102 | 0.000 |
| **Recall** | 0.858 | 0.858 | 0.858 | 0.827 | 0.773 | 0.693 | 0.453 | 0.098 | 0.000 |

| Threshold $\tau$ | 0.10 | 0.20 | 0.30 | 0.40 | 0.50 | 0.60 | 0.70 | 0.80 | 0.90 |
| --- | --- | --- | --- | --- | --- | --- | --- | --- | --- |
| 72 hpf |  |  |  |  |  |  |  |  |  |
| **Accuracy** | 0.737 | 0.732 | 0.726 | 0.689 | 0.635 | 0.578 | 0.460 | 0.236 | 0.034 |
| **Precision** | 0.945 | 0.941 | 0.937 | 0.909 | 0.865 | 0.816 | 0.702 | 0.425 | 0.072 |
| **Recall** | 0.769 | 0.767 | 0.764 | 0.740 | 0.705 | 0.665 | 0.571 | 0.346 | 0.059 |

#### Feature extraction

StarDist-3D provided as an output 32-bit float tif files that were converted to integer data type. Each segmented nucleus was identified by an integer value and all segmented nuclei touching the borders of the image were excluded from the analysis. The images were rescaled to obtain almost isovolumetric voxels that would facilitate the extraction of some of the features of interests. The script iterated through all nuclei in the image to measure the volume, the longer, intermediate and shorter axes from the centroid, the primary and secondary nematic axes and the centroid coordinates. The primary nematic was calculated as the longest pairwise distance between points composing the nuclear boundaries. The secondary nematic was defined as the maximum pairwise distance between points of the nuclear boundaries lying on the midplane to the primary nematic. Since nuclei supposedly do not touch each other, the nuclear boundaries of each nucleus were dilated to count the number of touching neighbors at different radii. To computer the neighbour statistics of each nucleus, a touch matrix was generated, reporting the centroid-to-centroid distance for each touching neighbour. In this way, the touching and proximal neighbours and the internuclear distances were computed for each nucleus in the ROI.

#### Image smoothing of extracted features

To explore the local spatial variations of extracted features, an automatic script was written to scan each image StarDist-3D prediction with a 3-dimensional window of a given size (Fig. S2 B, C). The images shown in the main figures were obtained with a 10×10×10 *µ*m size window. The mean value for the feature of choice was measured within the volumetric window, e.g. the average number of contacts. Since the volumetric windows were overlapping to by 20%, each pixel was averaged by the times the window looped on it. This created a smoothed image stack where each pixel reported the mean value for the measurement of interest. This analysis was performed using Python3.9.

#### Selection of the region of interests

Nuclei within regions of interest (ROI) were manually selected using a Python script that enables the visualization and user interaction with the dataset using Napari (v0.4.18). In this way, the user can draw a polygon over the area of interest and specify the range of slices desired. The script, then, iterates through all the labelled nuclei present within this region and creates a new image stack containing only those nuclei. Then, another Napari GUI automatically opens to allow the user to manually mark the labeled objects that were erroneously included in the new image. In this way, it was possible to correct the image and obtain a csv file reporting the labels of nuclei contained in the selected ROI.

### Data analysis

#### Analysis of the orientational order of nuclei

The description of a nematic liquid crystals involves the quantification of the order in the system. The average angle between the primary nematic axis of each nucleus and the primary nematics of its touching neighbors was calculated. The calculated averaged angles were used to compute the scalar order parameter S in selected ROIs:

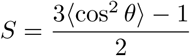

where *θ* is the angle between the liquid-crystal molecular axis and the local director. Note that for a completely random and isotropic sample, S = 0, whereas for a perfectly aligned sample S=1. For a typical nematic liquid crystal sample, 0.4 *<* S *<* 0.9. The same procedure was performed for the secondary nematic axes.

#### Analysis of the radial distribution function

The radial distribution function *g*(*r*) provides a statistical description of the local packing and particle density of a system. It is defined as:

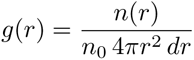

where *n*(*r*) is the number of particles in a spherical shell of radius *r* and thickness *dr*, *n*_0_ is the average number density of particles and 4*πr*^2^ *dr* is the volume of the spherical shell. The *g*(*r*) was calculated iteratively for the increment *dr* steadily increasing the radius of the external spherical shell, with a maximal radius of 50 *µ*m. To calculate the number density of nuclei within the ROI, the nuclear boundaries were dilated so that the space between the nuclei would be completely filled. Then, this labelled object was eroded to mask the nuclei in the ROI. The number density was calculated as the ratio between the number of nuclei in the mask and the volume of the mask itself. No compensation for edge effects was performed.

#### Data visualization and statistics

All statistical tests used are indicated in the figure or table legends, as well as the definitions of error bars. Likewise, p values and sample sizes are reported in the figure legends or in dedicated tables. After visual inspection of nuclei in the selected ROIs, a lower and upper threshold of nuclear volumes was used to filter the data obtained after extraction of all features of interests from the StarDist-3D predictions. More specifically, we used a lower threshold of 50 *µ*m^3^ and upper thresholds of 400 *µ*m^3^ for nuclei in embryos staged between 24 and 48 hpf in the retina. For later stages, we used a lower threshold of of 20 *µ*m^3^. This was done to reduce the number of erroneously segmented nuclei. Data visualization and statistical analysis were performed using the Matplotlib, Seaborn and Scipy Statistics packages in Python 3.9. Further information about the exact libraries, together with their versions, are detailed in the Github repository.

## Data and code availability

Imaging and model data will be made available on the dedicated Github repository and on the BioImage Model Zoo platform.

## Acknowledgements

We thank the Cell Biology of Tissue Morphogenesis laboratory for fruitful project and manuscript discussion. Nerli E. is thanked for experimental help. Weigert M. is thanked for his precious help in setting up StarDist-3D for this study. We thank Nerli E., Ramos A.P., Matejcic M., Petridou N., Haase P. and Grill S. for thoughtful feedback on the manuscript. We are grateful to the Fish Facility and the Imaging Facility of the Gulbenkian Institute for Molecular Medicine (GIMM) for experimental support. L.C.F. was a member of the Integrative Biology and Biomedicine PhD program and was supported by the Fundação para a Cîencia e a Tecnologia PhD studentship (2020.06407.BD). C.N. was supported by the Fundação Calouste Gulbenkian-IGC, by the European Research Council (ERC) under the European Union’s Horizon 2020 research and innovation program (no. 819046), and by the Fundação para a Cîencia e a Tecnologia Investigator grant (CEECIND/03268/2018). C.D.M. and A.Q.R. acknowledge funding from the Max Planck Society through the MPI-CBG. A.Q.R. additionally acknowledges funding support through the ELBE Postdoctoral Fellowship program. R.H. acknowledges support by the Deutsche Forschungsgemeinschaft under Germany’s Excellence Strategy—EXC2068–Cluster of Excellence Physics of Life of TU Dresden and by the programme Center of Excellence for AI-research “Center for Scalable Data Analytics and Artificial Intelligence Dresden/Leipzig”.

## Authors contributions

C.N., C.D.M. and L.C.F. conceptualized work plan and methodology. L.C.F. performed the majority of the experiments and analysis. A.Q.R. and R.H. helped with the image analysis pipeline. C.D.M. conceptualized and performed the theoretical modelling. C.N. and L.C.F. wrote the manuscript with the help of all the other authors.

## Declarations

The authors declare no competing financial interests.

## Supplementary section

**Fig. S1.**
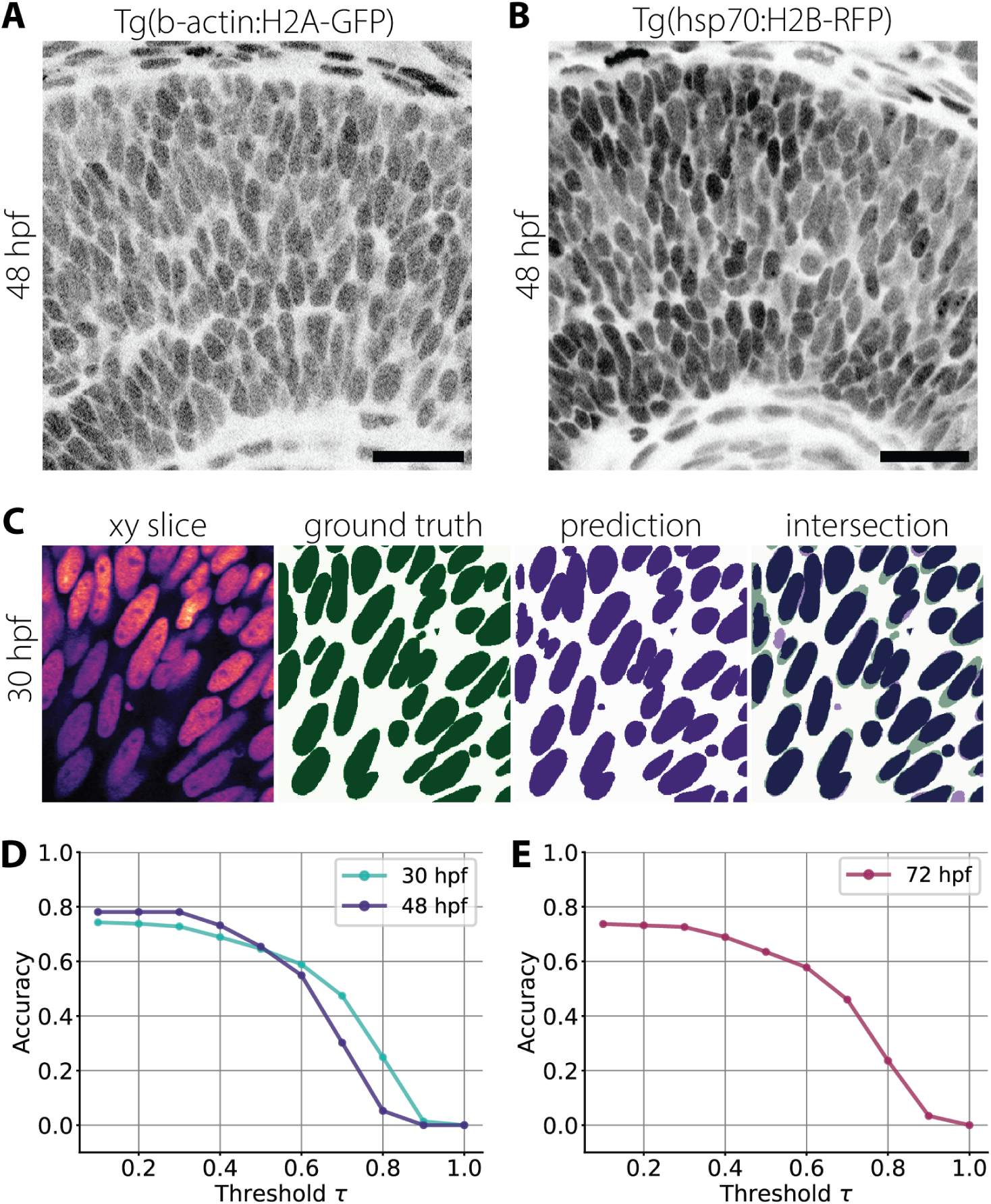
Training and evaluation of custom StarDist-3D models. **A), B)** Representative confocal sections of 48 hpf retinas from Tg(hsp70:H2B-RFP) in panel (A) and Tg(b-act:H2A-GFP) embryos in panel (B). Scale bar is 20 *µ*m. **C)** Representative optical slices of one cropped stack from the test dataset. Corresponding manually annotated ground truth image and StarDist-3D prediction are shown, together with their overlap. **D)** Accuracy for several IoU thresholds for evaluation datasets from 30 hpf and 48 hpf retinas for StarDist-3D model A. The number of test ground-truth stacks per stage is: 30 hpf, N = 2 images, n = 272 nuclei; 48 hpf, N = 3; n = 225. **E)** Accuracy for several IoU thresholds for evaluation datasets from 72 hpf retinas for StarDist-3D model B. 72 hpf, N images = 4, n nuclei = 647.

**Fig. S2.**
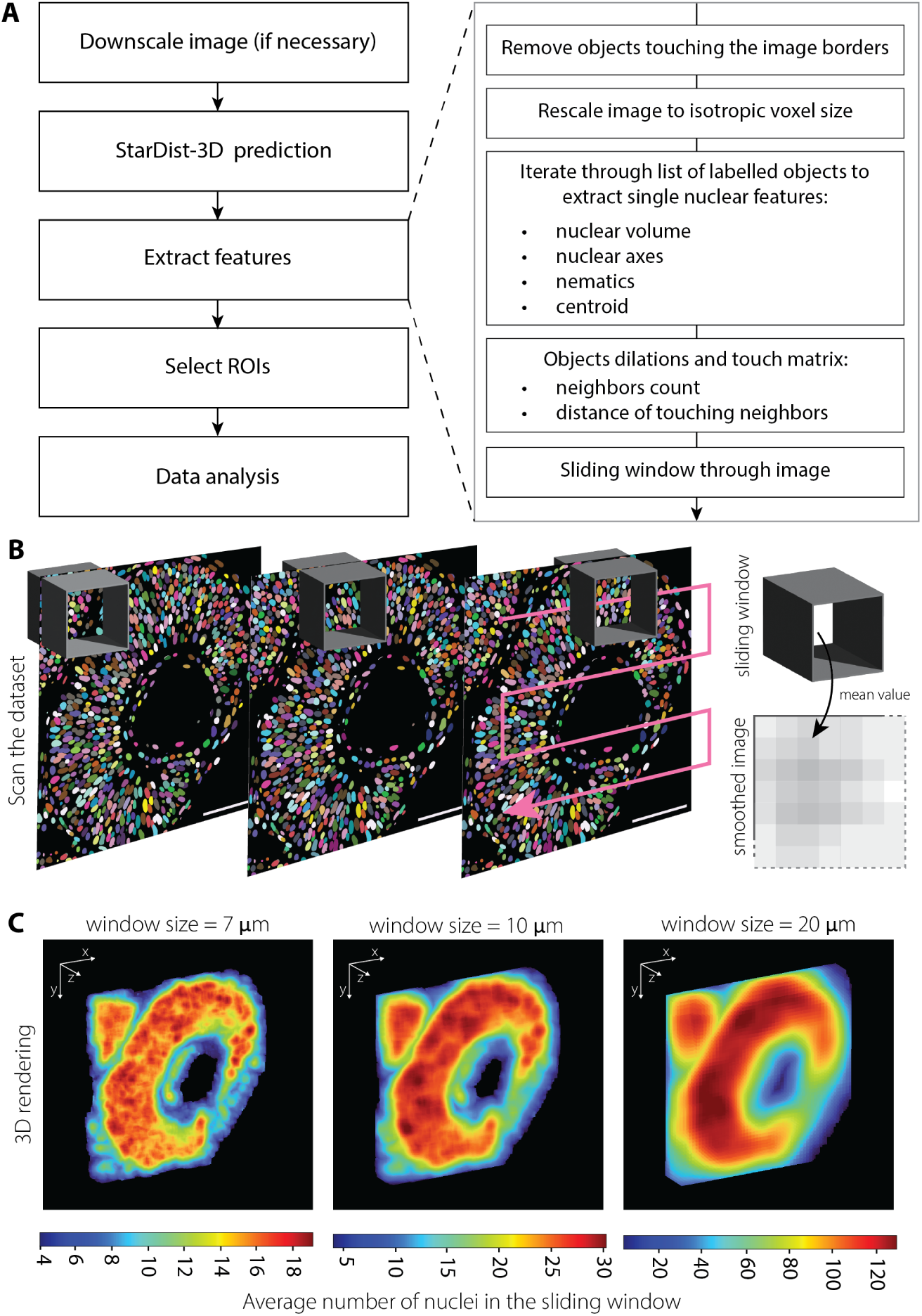
Analysis pipeline of volumetric imaging dataset. **A)** Workflow of the established 3D image analysis pipeline. **B)** Schematics showing the procedure to smooth volumetric imaging dataset. The cubic window iterated through the image stack of a 42 hpf segmented retina and the average value for the feature of interest was calculated and assigned to the pixels inside the window. Ultimately, a smoothed dataset reporting average values computed for each pixel is generated. Scale bar is 50 *µ*m. **C)** Representative 3D renderings of the 42 hpf segmented retina shown in panel (B) after smoothing the number of nuclei within different window sizes. The pixel intensities correspond to the average number of nuclei within the window.

**Fig. S3.**
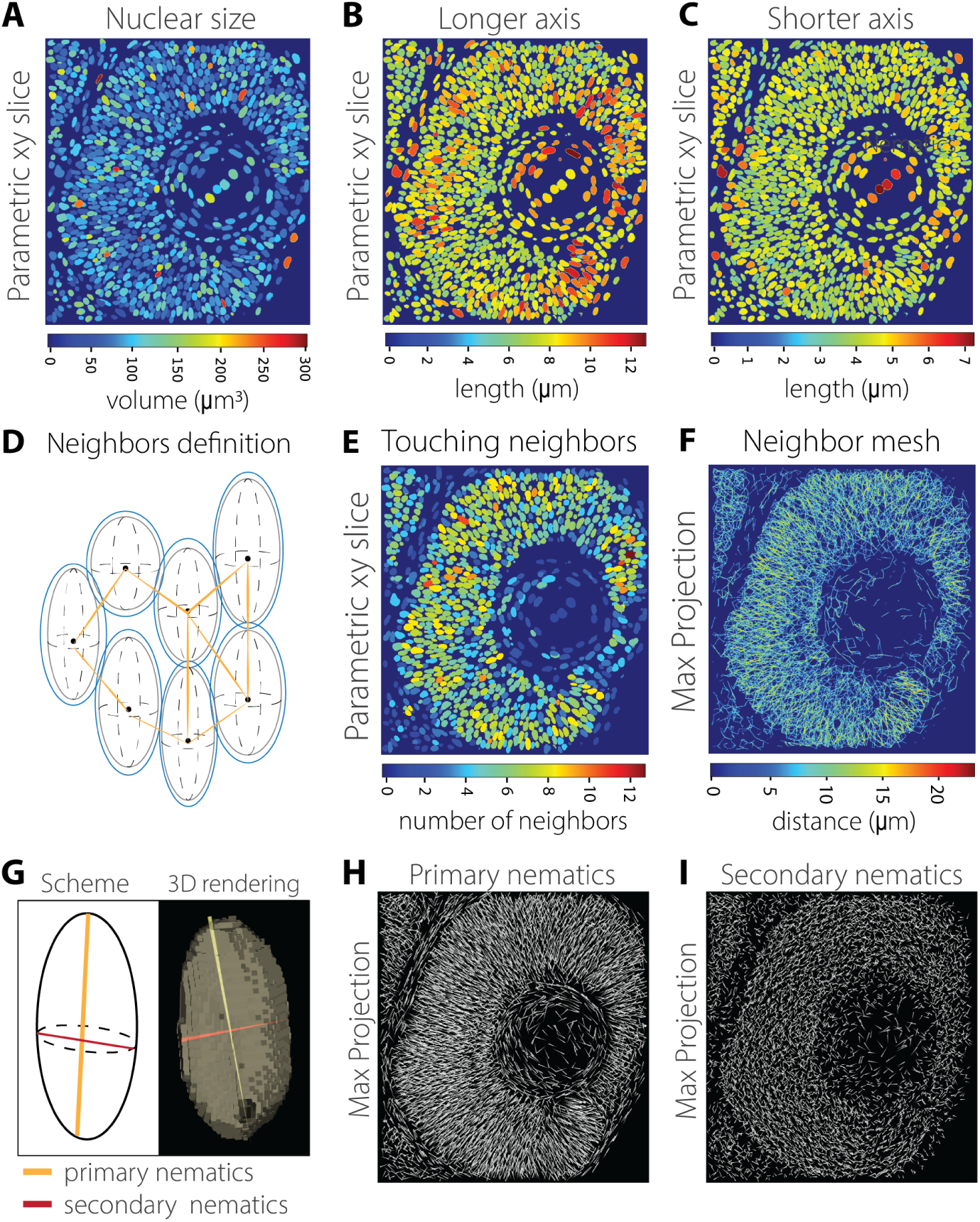
Extraction of nuclear shape and neighbourhood descriptors. **A), B), C), E)** Representatives optical slices of parametric images of a 42 hpf segmented retina. Parametric images showing volume of nuclei (A), lengths of the longer (B) and shorter (C) axes and number of touching neighbours (E). **D)** Schematic showing the definition of touching neighbours. Boundaries of the nuclei were expanded for a given radius (dilated nuclei outlined in blue) and internuclear distance between touching neighbours was computed to form a mesh (in orange). **F)** Maximum intensity projection of distance mesh between touching nuclei from panel (E). **G)** Schematics of primary and secondary nematic axes. **H), I)** Maximum intensity projections of primary (H) and secondary (I) nematics in 42 hpf retina.

**Fig. S4.**
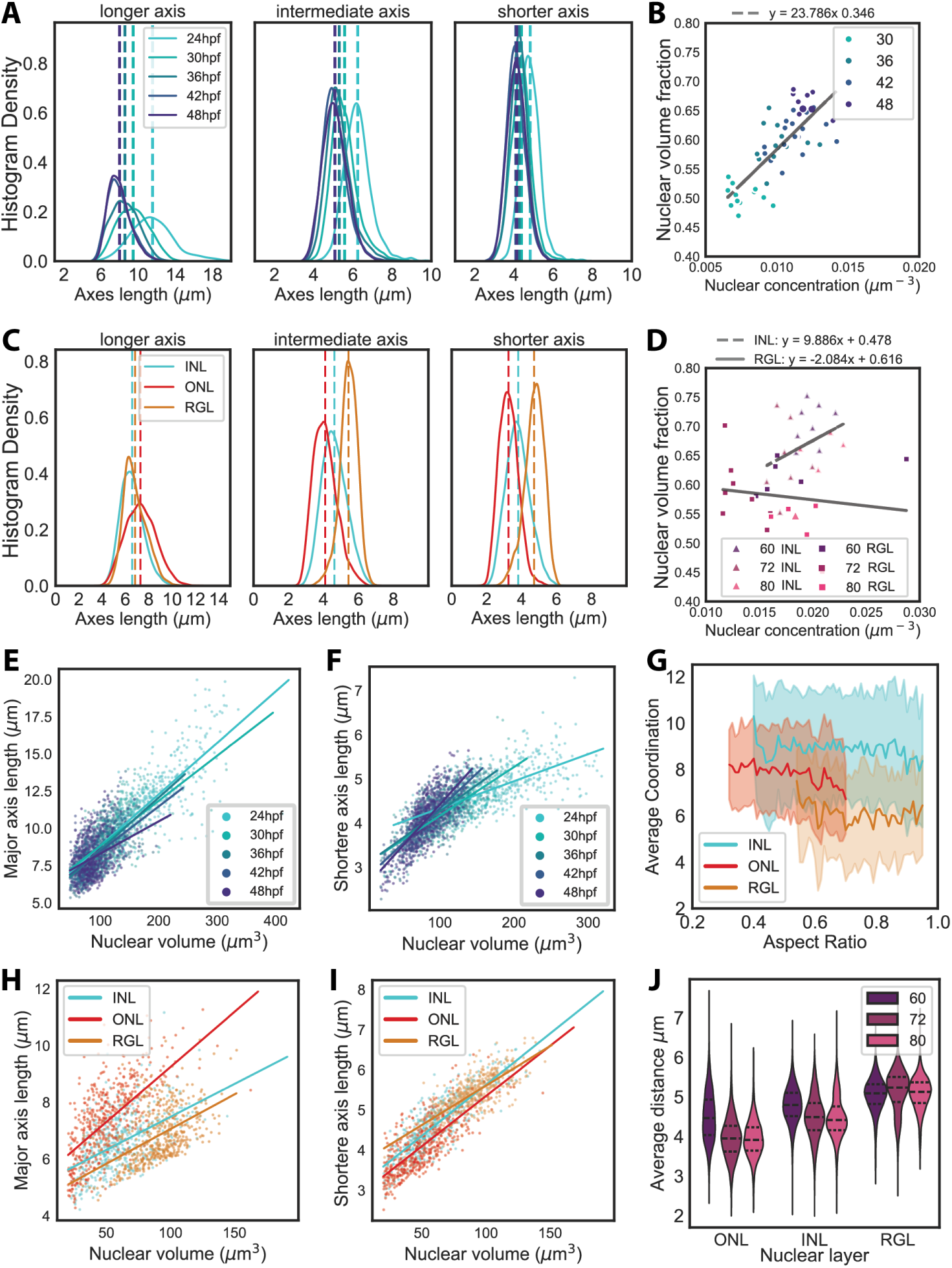
Nuclear shape and positioning changes during the transition from a PSE to a laminated retina. **A), C)** Histogram density distributions of nuclear axis lengths of nuclei in the RNE during the proliferative phase (A) and in 72 hpf retinas (C). Color-coded and segmented lines in panels (A, C) indicate the mean values for each distribution. **B), D)** Scatter plots showing linear correlation between nuclear volume fraction and concentration during the proliferative phase of the RNE. **C)** Histogram density distributions of nuclear axis lengths in 72 hpf retinas. Color-coded and segmented lines in panels (A, C) indicate the mean values for each distribution. **E), F)** Scatter plot between nuclear volume and longer axis lengths (E) and shorter axis lengths (F) during the proliferative phase of the RNE. Pearson’s correlation coefficient for panel (E): 24 hpf, r = 0.79; 30 hpf, r = 0.65; 36 hpf, r = 0.66; 42 hpf, r = 0.56; 48 hpf, r = 0.36. Pearson’s correlation coefficient for panel (F): 24 hpf, r = 0.64; 30 hpf, r = 0.72; 36 hpf, r = 0.71; 42 hpf, r = 0.75; 48 hpf, r = 0.75. **G)** Correlation between average coordination and nuclear aspect ratios in 72 hpf retinas across the three nuclear layers. Solid lines show mean values, shaded area indicates the standard deviation from the mean. **H), I)** Scatter plot between nuclear volume and longer axis lengths (H) and shorter axis lengths (I) in 72 hpf retinas. Pearson’s correlation coefficient for panel (H): ONL, r = 0.00; INL, r = 0.00; RGL, r = 0.00. Pearson’s correlation coefficient for panel (I): ONL, r = 0.00; INL, r = 0.00; RGL, r = 0.00. **J)** Distributions of average distances across the three nuclear layers between 60 and 80 hpf.

**Fig. S5.**
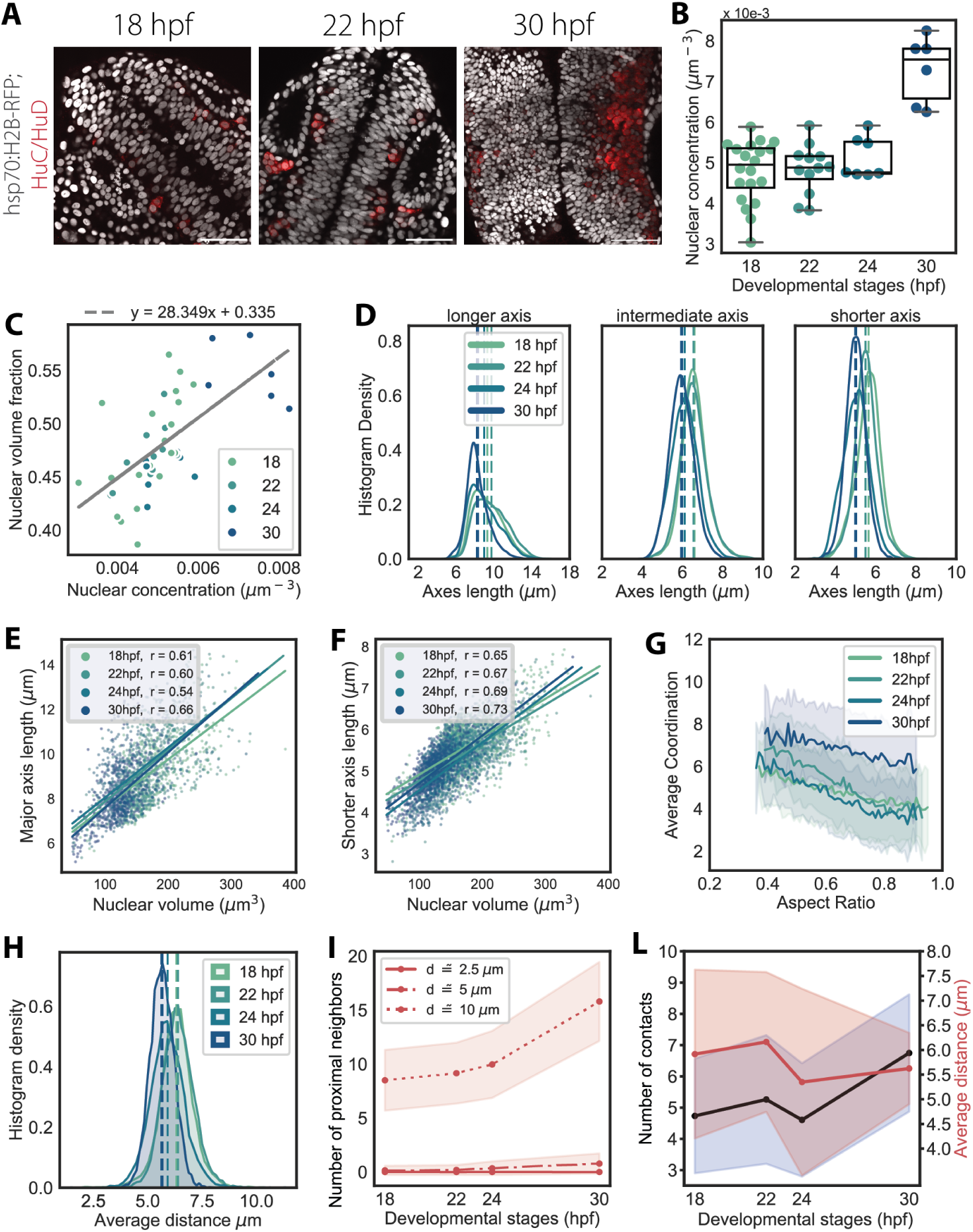
Single-nuclear shape descriptors to follow nuclear packing changes in the hindbrain. **A)** Neuronal differentiation starts around 18 hpf in the hindbrain. Huc/HuD-positive neurons are marked in red. Nuclei are labelled with Tg(hsp70:H2B-RFP). Scale bar is 50 *µ*m. **B)** Concentration of nuclei found within the ROIs of Fig. 3 C. **C)** Scatter plot showing linear correlation between nuclear volume fraction and concentration during early stages of hindbrain development. D) Histogram density distributions of nuclear axis lengths over development. Segmented lines indicate the mean values for each distribution. **E), F)** Scatter plot between nuclear volume and longer axis lengths (E) and shorter axis lengths (F) to detect their correlation. Pearson’s correlation coefficient (r) for panel (E): 18 hpf, r = 0.61; 22 hpf, r = 0.60; 24 hpf, r = 0.54; 30 hpf, r = 0.66. Pearson’s correlation coefficient (r) for panel (F): 18 hpf, r = 0.65; 22 hpf, r = 0.67; 24 hpf, r = 0.69; 30 hpf, r = 0.73. **G)** Correlation between average coordination and nuclear aspect ratios. Solid lines show mean values, shaded area indicate the standard deviation from the mean. H) Histogram density distributions of average distances between touching neighbours. **I)** Number of proximal neighbours within different distance radii. **J)** Mean number of contacts within ROIs increase, while the average distance between touching neighbours oscillates around the same value. Solid lines report mean values, shaded area indicate the standard deviation from the mean.

**Fig. S6.**
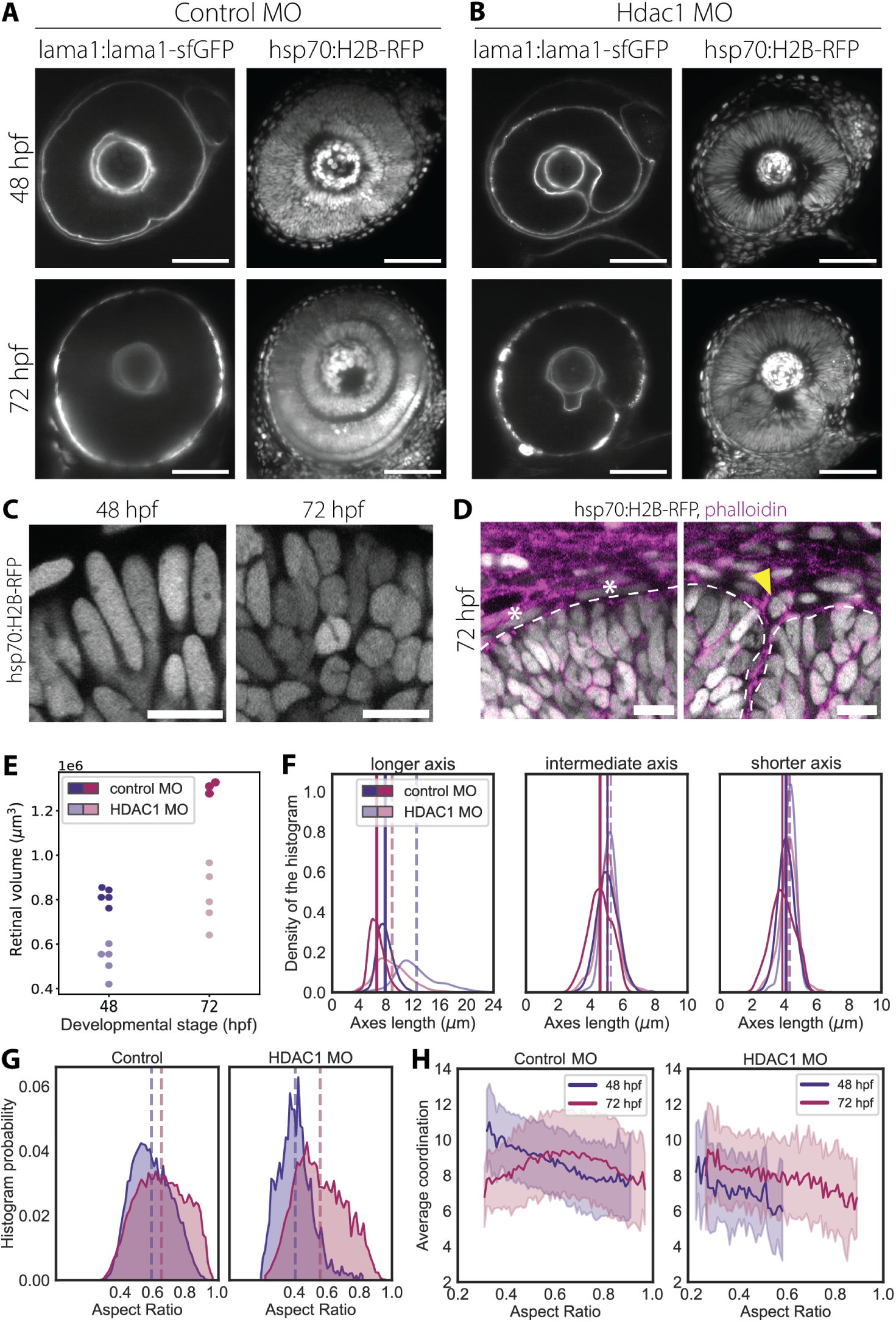
HDAC1 morphants feature reduced tissue growth and deformed nuclear shapes. **A), B)** Representative confocal sections of the retina of 48 and 72 hpf zebrafish embryos that were injected either with control morpholino (MO) (panel A) or with HDAC1 MO (panel B). Nuclei are labelled with tg(hsp70:H2B-RFP), the basal lamina is marked with tg(lama1:lama1-sfGFP). Scale bar is 50 *µ*m. **C)** Representative cropped confocal sections of 48 hpf and 72 hpf retinas in HDAC1 morphants. Nuclei are labelled with tg(hsp70:H2B-RFP). Scale bar is 10 *µ*m. **D)** Representative cropped confocal sections of different regions of 72 hpf retina from HDAC1 morphant. The sections show part of the RNE and RPE in regions where the tissue is not buckled (left) and where it is buckled (right). Nuclei of RPE cells appear to be stretched (asterisks) where the tissue is not buckled, while they are more rounded over buckled regions (triangle). Nuclei are labelled with tg(hsp70:H2B-RFP) in grey; the actomyosin skeleton is labelled with phalloidin in magenta. Scale bar is 10 *µ*m. **E)** Eye volume from quantified from tg(lama1:lama1-sfGFP) embryos. N = 3 to 5 embryos per condition. **F)** Histogram density distributions of nuclear axis lengths of nuclei in control and HDAC1 morphants at 48 hpf and 72 hpf. Color-coded and segmented lines indicate the mean values distribution of HDAC1 morphants. Color-coded and solid lines indicate the mean values distribution of control morphants. **G)**Histogram density distributions of nuclear aspect ratios, which were defined as the ratio between the longer axis length and the mean length between the intermediate and shorter axes. **H)** Correlation between average coordination and nuclear aspect ratios in 48 hpf and 72 hpf retinas from control (left) and HDAC1 (right) morphants. Solid lines show mean values, shaded area indicate the standard deviation from the mean.

## Nuclear Packing Buckling Instability Theory Supplement

### 1 Introduction and Set-Up

In order to model the effect that the nuclear packing environment in the optic cup has on tissue stability and the potential for a buckling transition, we take inspiration from Trushko et al. (2020) [1]. There, Trushko and coauthors render a leading-order continuum model of an epithelium growing under spherical confinement as an (initially) circular elastic ring confined in a circular geometry of radius *R*. When the elastic ring becomes larger than the size of the confining circle, a buckling instability sets in and the elastic ring breaks circular symmetry and buckles. More in depth treatments of this problem have also been carried out to higher order, for example by Napoli and Turzi [2], although the dominant, leading order behavior is unchanged. Here, we choose to follow the simpler approach of Trushko et al., but additionally capture the nuclear packing environment in the coarse-grained continuum model as two distinct material states, with material properties dependent in a switch-like manner on the packing fraction.

We begin by adapting the global energy per unit length of the confined 1d elastica ring, for *L* the current length of the elastica and *L*_0_ the equilibrium (preferred) length without any constraints:

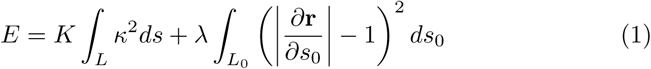

where we chose here to ignore any potential energetic contribution from distortions to the confining medium and assume instead that the combination of ECM and other tissues around the optic cup are significantly stiffer and effectively incompressible. The first term captures the bending energy of the elastica, while the second term captures any contribution from compression (or extension) along the tissue.

In order to derive a critical strain for the onset of the buckling instability and investigate how a two-state system of internal stiffnesses in the tissue affects this instability, we will need a simple, minimal buckled elastica shape (confined within a circle) to compare with the elastica remaining circular at the cost of compression energy. As Trushko et al. do, we also choose to use a simple trigonometric representation of the initially buckled shape:

**Fig. TS1:**
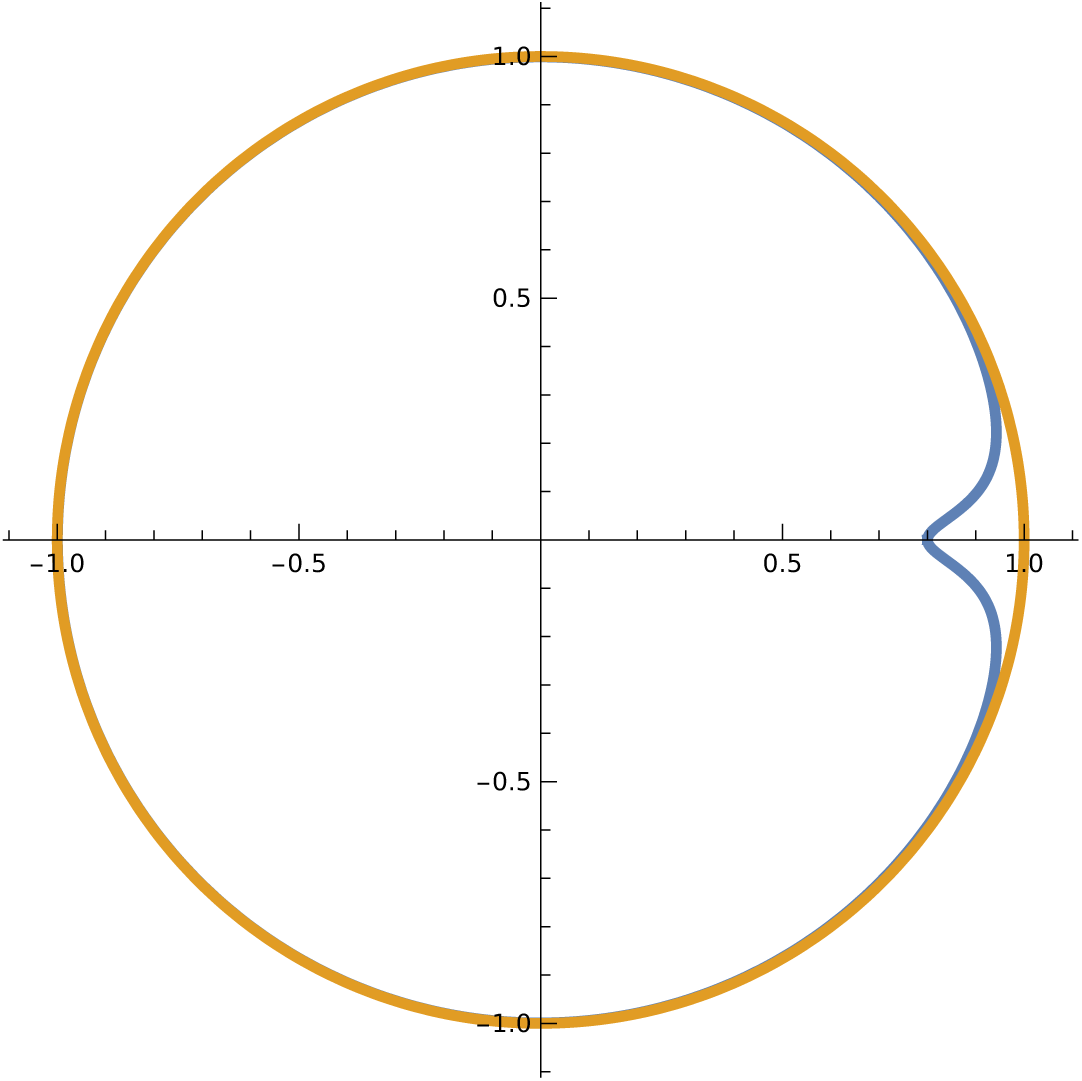
Example buckled configuration, shown here for an elastica (blue) constrained inside a rigid unit circle (orange), with *δ* = 0.2 and *α* = 0.05.

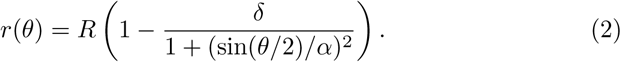

Unlike Trushko et al., though, we assume that the buckle occurs at *θ* = 0 and that there is only one buckling site, requiring the extra factor of 1*/*2 inside the argument of the sine function. Here, *α* and *δ* are dimensionless parameters controlling the shape of the buckled region, with *δ* setting the depth of the buckled fold and *α* setting the width of the buckled region. The length of an elastica parameterized in this way is given by:

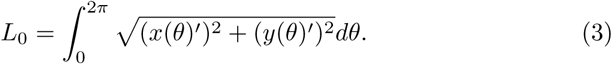

As we intend to compare a just-buckled elastica shape with its compressed, pre-buckled counterpart, we may assume that the amplitude and extent of the buckled region are small and we accordingly set *α, δ* ≪ 1. We may then expand in small quantities, allowing us the write, to leading order:

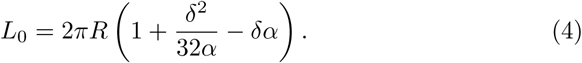

It is useful here to introduce the excess strain, Δ*ε* = (*L*_0_ - 2*πR*)*/*2*πR*, leading to:

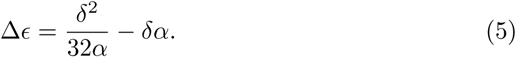

Continuing to follow Trushko et al., we treat the expression for the bending energy similarly, expanding to leading order in small quantities, and writing:

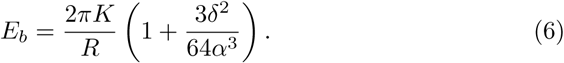

Note that naively taking our small quantities back to zero here is suddenly problematic as it would imply a diverging energy! This is an early hint that the scaling of *δ* and *α* are not independent of one another for a physically realistic buckled region, and that *δ* should scale faster than *α* in order to cut off the unphysical divergence. How to ensure that we are considering a physically realistic buckled region, then? We wish to work with the elastica shapes that, for a given excess strain Δ*ε*, minimize the bending energy. We can write the Lagrangian for this optimization as:

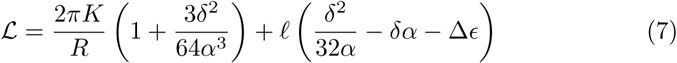

with *ℓ* the Lagrange multiplier enforcing the condition arising from the given excess strain. Computing the requisite gradients and solving for *δ* and *α* in terms of Δ*ε* finally yields the following scaling relations:

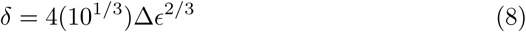

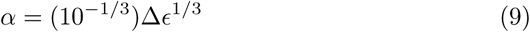

which may be then re-substituted back into the expression for the bending energy to obtain the scaling of the pure-bend configuration’s energy with Δ*∈*:

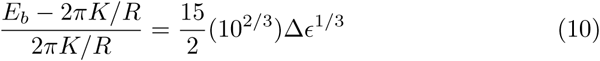

Meanwhile, the elastic compressive energy of the confined elastica is simply quadratic in the excess strain, as it would be for a spring or any other Hookean material, and the pure-compression configuration’s energy scales with Δ*∈* as:

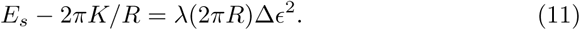

Initially, for very small values of Δ*ε*, the pure-compression configuration is less energetically costly and, despite the fact that there is some excess strain, the elastica does not buckle to adopt curvature. At larger values of Δ*ε*, however, the pure-bend configuration is favored over pure compression and buckling would therefore be expected to take place. Since the energetic costs of the two possible configurations switch places in relative costliness as Δ*ε* increases, there must be a critical value of the excess strain, Δ*∈_c_*, where the two energies are equal. This Δ*∈_c_* therefore captures the onset of the buckling instability and can be found by setting *E_b_* = *E_s_*:

**Fig. TS2:**
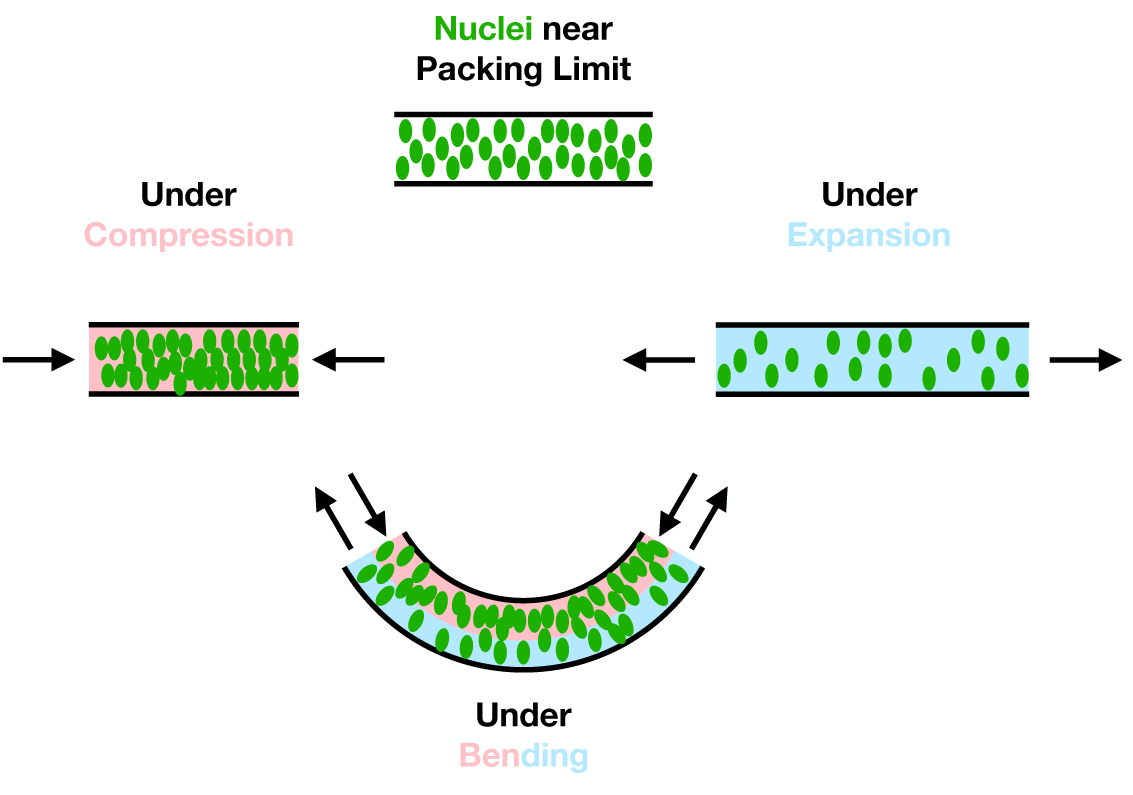
Schematic indicating the interplay of the internal packing state of the tissue’s nuclei with externally imposed deformations: compression, expansion, and bending. Compression can drive the mechanics from cellularly dominated to nuclearly dominated while expansion is the reverse. Bending creates a mix of both.

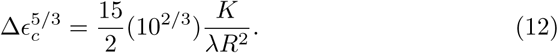

Note that for a fixed size of the constraining environment, the critical excess strain required to initiate buckling is thus controlled entirely by (*K/λ*)^3^*^/^*^5^. We have now gathered the necessary ingredients to consider the two-state-stiffness tissue we propose as an analogy for the developing RNE.

### 2 Nuclear jamming and the onset of buckling

To this point we have closely followed the approach of Trushko et al. in deriving the scaling of the energies and the critical buckling excess strain for the constrained elastica. In order to incorporate multiple internal states into the elastica that depend on Δ*ε* and may alter its stiffness, however, now requires us to set off in a new direction. We wish to define new compression and bending moduli, *λ* and *K* that are functions of the preferred length, *L*_0_, of the elastica in a way that captures the onset of nuclear jamming in the RNE and the attendant switch from *cellular* -associated stiffness in the unjammed state to *nuclear* -associated stiffness in the jammed state. We write:

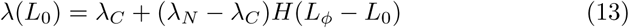

for *λ_C_* the cellular compression stiffness, *λ_N_* the nuclear compression stiffness, *L_ϕ_* the tissue length below which jamming occurs given the number of nuclei present, and *H*() the Heaviside step function. We further assume that *λ_N_ > λ_C_*, as the nuclear envelope is commonly understood to be the stiffest component of the cell, often in the range of 2-10 times stiffer than the rest of the cell, and in many cases one might even expect *λ_N_* ≫ *λ_C_* [3, 4]. We also introduce an *L*_0_-dependence to the bending rigidity in a similar manner:

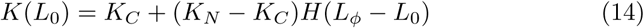

for *K_C_* and *K_N_* the cellular and nuclear-associated bending rigidities, respectively. Recall that the bending rigidity generically scales with the compression/stretching stiffness (i.e. the Young’s modulus), *K ~ λ*, for simple homogeneous isotropic materials. This is because one can think of the source of bending energy for a downward directed bend as a profile of stretch above the mid-line (or mid-plane for a bending plate or cantilever) and compression below. In the regime of our tissue dominated by the cellular mechanics, where the nuclei are not densely packed enough to begin mechanically interacting, this same simple scaling holds and we may write *K_C_ ∼ λ_C_*. However, something more interesting happens in the close-packed, nuclear-jammed state. Here, under a bending deformation, the tissue on the side of the mid-line experiencing compression remains in the regime controlled by the nuclear mechanics, but on the opposite side, where the tissue is stretched and expanded, the nuclei fall out of their jammed state and cellular mechanics dominate. The scaling of *K_N_* is therefore more subtle, and can be realized as *K_N_* ∼ (*λ_C_* + *λ_N_*)*/*2 so long as the system is not too deep into the jammed regime even at relaxed length, since exactly half the tissue experiences one compression/stretching stiffness and the other half experiences the other. For convenience we employ a scale factor, *k* = *h*^3^*/*12(1 − *ν*^2^), for *h* the tissue height and *ν* its Poisson ratio in order to capture the rest of the bending rigidity [5] and write:

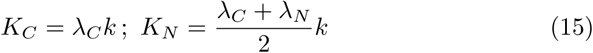

Let us now consider the different possible relevant regimes of *L*_0_, 2*πR*, and *L_ϕ_*. We may begin by enforcing the assumption that *L*_0_ *>* 2*πR* as otherwise there is no longer any effective confinement and nothing interesting happens. Thus there are three remaining regimes to investigate: (1) *L_ϕ_ <* 2*πR < L*_0_, (2) 2*πR < L_ϕ_ < L*_0_, and (3) 2*πR < L*_0_ *< L_ϕ_*.

In the first regime, all the other length scales in the system are greater than the length below which jamming and the attendant nuclear-associated stiffnesses set in and the tissue is solidly in the cellular mechanics dominated state. The critical buckling excess strain is therefore given by:

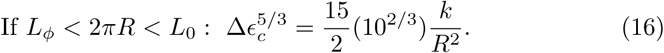

In the third regime, on the other hand, the other length scales in the system are below the jamming threshold length, *L_ϕ_* and thus the tissue is solidly in the nuclear mechanics dominated state. Now, the critical buckling excess strain is instead given by:

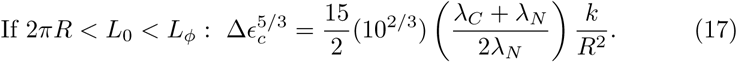

Note that the critical excess strain has decreased here relative to its value in the first regime, perhaps significantly so if *λ_N_ ≫ λ_C_*, meaning that buckling becomes easier to initiate once the tissue is (nuclearly) jammed.

What of the second, intermediate regime? Here the pure-bend configuration has tissue length *L*_0_ *> L_ϕ_* and resides in the cellular mechanics dominated state, while the pure-compression configuration has tissue length 2*πR < L_ϕ_* and resides in the nuclear mechanics dominated state. However, the compressional energy we used earlier to derive Δ*σ_c_* must be adjusted in this case, as compressing the tissue from its preferred length *L*_0_ down to the current, unbuckled length 2*πR* requires crossing the *L_ϕ_* boundary and switching stiffness regimes. In other words, some of the work done compressing the tissue is done in the cellular mechanics dominated state, even if the tissue is currently jammed in the nuclear mechanics dominated state. The corrected stretch/compression energy now reads:

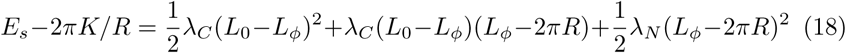

or, rearranging and re-writing the per-unit-length energy in terms of Δ*∈* gives:

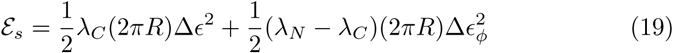

where Δ*∈_ϕ_* is the excess strain to the jamming transition boundary, Δ*∈_ϕ_* = (*L_ϕ_* − 2*πR*)*/*2*πR*. We can now use this new energy to derive the adjusted critical excess strain in this crossover regime, finding:

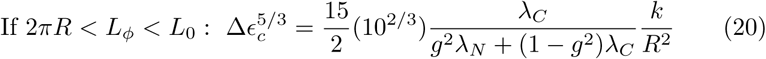

for *g* = Δ*∈_ϕ_/*Δ*∈*. Note that 0 *< g <* 1, as Δ*∈_ϕ_* ranges from 0 to Δ*∈* as *L_ϕ_* ranges from 2*πR* to *L*_0_. Note further that, in this regime, as *g* → 1, 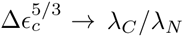 and not *K_C_/λ_N_*, leaving a discontinuous jump in the behavior of the critical exponent, as can be seen in Fig. TS3. A more complex, quasi-2D shell treatment of the problem that includes the varying amounts of extension and compression due to bend away from the mid-line of the elastica for different thicknesses would be expected to resolve this discontinuity by introducing a term coupling the stretch and the bend, but would also be expected to retain the same qualitative feature of the critical exponent diving down below the value for nuclear-mechanics dominated regime then sharply recovering to match that value very close to Δ*∈_ϕ_* = Δ*∈*. Such a shell theoretic treatment of a system with strain-dependent two-state material properties is beyond the scope of what is required to understand the optic cup buckling susceptibility here, but nevertheless constitutes an interesting direction for future theoretical work.

**Fig. TS3:**
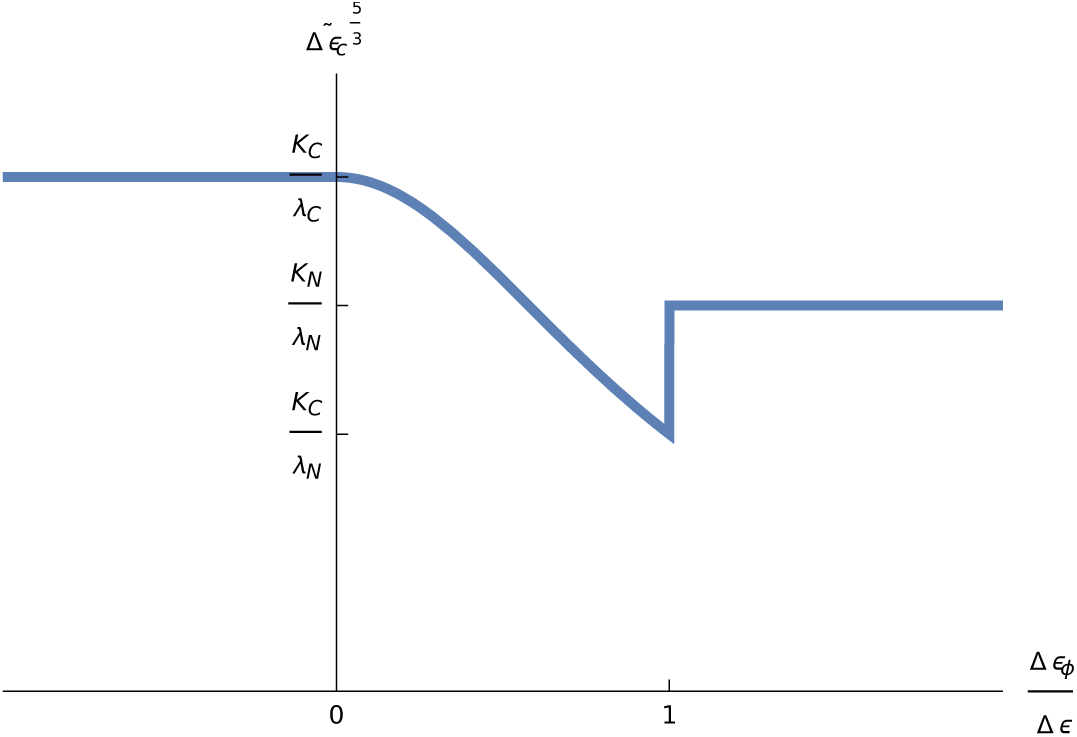
The normalized critical strain, 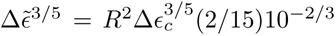 as a function of *g*. Note that the critical strain is relatively high in the regime dominated by cellular mechanics and relatively low in the regime dominated by nuclear mechanics. In the crossover region, on approach to the nuclear-mechanics dominated regime, the onset of a buckling instability becomes even easier still, with a decrease of 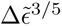 by a factor of 10, for examplee, if *λ_N_* ∼ 10*λ_C_*, which is in the physiologically relevant range [3, 4].

Returning to the case of the developing hemispheric pseudostratified retinal neuroepithelium, the fact that the approach to a jammed, nuclear-mechanics dominated internal state is accompanied by a sharply decreased critical buckling strain presents a harsh mechanical constraint on the proliferation and arrangement of the RNE as the optic cup develops. Indeed, if the developmental program wishes to ensure that the shape of the optic cup is preserved – as it must be in order to guarantee proper function of the eye – then the final approach to the nuclear-mechanics dominated internal state is to be avoided. Strikingly, in the wildtype optic cup, it is precisely when the nuclear packings begin to approach limiting values that neurogenesis and lamination begin, and the nuclear arrangements are allowed to adopt spatial order in addition to their pre-existing orientational order. These more crystalline packing states are associated with significantly higher free volumes than in the disordered cases, opening up more room for the nuclei at the same densities, and allowing the tissue to remain in the cellular mechanics dominated regime, relatively safe from the onset of buckling instabilities. This analysis therefore predicts that, if the transition to neurogenesis and lamination is interfered with, then the growing optic cup may no longer be able to avoid the marked decrease in the critical buckling strain, and uncontrolled, ectopic buckles and folds would very likely result, matching what we observe in the HDAC mutants.

## References

[1] Smart, I. H. Proliferative characteristics of the ependymal layer during the early development of the spinal cord in the mouse (1972).

[2] Miyata, T. Development of three-dimensional architecture of the neuroepithelium: Role of pseudostratification and cellular ‘community’ 50 (2008). URL https://onlinelibrary.wiley.com/doi/10.1111/j.1440-169X.2007.00980.x.

[3] Ferreira, M. A., Despin-Guitard, E., Duarte, F., Degond, P. & Theveneau, E. Interkinetic nuclear movements promote apical expansion in pseudostratified epithelia at the expense of apicobasal elongation 15, 1–24.

[4] Guerrero, P. et al. Neuronal differentiation influences progenitor arrangement in the vertebrate neuroepithelium 146.

[5] Kicheva, A. et al. Coordination of progenitor specification and growth in mouse and chick spinal cord 345.

[6] Iulianella, A., Sharma, M., Durnin, M., Vanden Heuvel, G. B. & Trainor, P. A. Cux2 (Cutl2) integrates neural progenitor development with cell-cycle progression during spinal cord neurogenesis 135, 729–741. URL https://journals.biologists.com/dev/article/135/4/729/64918/Cux2-Cutl2-integrates-neural-progenitor.

[7] Bystron, I., Blakemore, C. & Rakic, P. Development of the human cerebral cortex: Boulder committee revisited 9, 110–122. URL https://www.nature.com/articles/nrn2252.

[8] Eiraku, M. et al. Self-organizing optic-cup morphogenesis in three-dimensional culture 472, 51–58. Publisher: Nature Publishing Group.

[9] Nakano, T. et al. Self-formation of optic cups and storable stratified neural retina from human ESCs 10, 771–785. URL 10.1016/j.stem.2012.05.009. Publisher: Elsevier Inc.

[10] Lancaster, M. A. et al. Cerebral organoids model human brain development and microcephaly 501, 373–379. URL https://www.nature.com/articles/nature12517.

[11] Meyer, E. J., Ikmi, A. & Gibson, M. C. Interkinetic nuclear migration is a broadly conserved feature of cell division in pseudostratified epithelia 21, 485–491. URL 10.1016/j.cub.2011.02.002. Publisher: Elsevier Ltd.

[12] Strzyz, P. J., Matejcic, M. & Norden, C. Heterogeneity, Cell Biology and Tissue Mechanics of Pseudostratified Epithelia: Coordination of Cell Divisions and Growth in Tightly Packed Tissues Vol. 325 (Elsevier Inc.). URL 10.1016/bs.ircmb.2016.02.004. Pages: 118 Publication Title: International Review of Cell and Molecular Biology.

[13] Norden, C. Pseudostratified epithelia – cell biology, diversity and roles in organ formation at a glance 130, 1859–1863.

[14] Bort, R., Signore, M., Tremblay, K., Barbera, J. P. M. & Zaret, K. S. Hex homeobox gene controls the transition of the endoderm to a pseudostratified, cell emergent epithelium for liver bud development 290, 44–56. URL https://linkinghub.elsevier.com/retrieve/pii/S0012160605007803.

[15] Gόmez, H. F., Dumond, M. S., Hodel, L., Vetter, R. & Iber, D. 3d cell neighbour dynamics in growing pseudostratified epithelia 10, 1–25.

[16] Schoenwolf, G. C. & Powers, M. L. Shaping of the chick neuroepithelium during primary and secondary neurulation: Role of cell elongation 218, 182–195. URL https://onlinelibrary.wiley.com/doi/10.1002/ar.1092180214.

[17] Sauer, F. C. Mitosis in the neural tube 62, 377–405. URL https://onlinelibrary.wiley.com/doi/10.1002/cne.900620207.

[18] Norden, C., Young, S., Link, B. A. & Harris, W. A. Actomyosin is the main driver of interkinetic nuclear migration in the retina 138, 1195–1208. URL 10.1016/j.cell.2009.06.032. Publisher: Elsevier Ltd.

[19] Leung, L., Klopper, A. V., Grill, S. W., Harris, W. A. & Norden, C. Apical migration of nuclei during g2 is a prerequisite for all nuclear motion in zebrafish neuroepithelia 139, 2635–2635. URL https://journals.biologists.com/dev/article/139/14/2635/45193/Apical-migration-of-nuclei-during-G2-is-a.

[20] Azizi, A. et al. Nuclear crowding and nonlinear diffusion during interkinetic nuclear migration in the zebrafish retina 9, 1–31.

[21] Mateǰcíc, M., Salbreux, G. & Norden, C. A non-cell-autonomous actin redistribution enables isotropic retinal growth 16, 1–29.

[22] Strzyz, P. J. et al. Interkinetic nuclear migration is centrosome independent and ensures apical cell division to maintain tissue integrity 32, 203–219. URL 10.1016/j.devcel.2014.12.001. Publisher: Elsevier Inc.

[23] Morin, X., Jaouen, F. & Durbec, P. Control of planar divisions by the g-protein regulator LGN maintains progenitors in the chick neuroepithelium 10, 1440–1448. URL https://www.nature.com/articles/nn1984.

[24] Nakajima, Y.-i., Meyer, E. J., Kroesen, A., McKinney, S. A. & Gibson, M. C. Epithelial junctions maintain tissue architecture by directing planar spindle orientation 500, 359–362. URL https://www.nature.com/articles/nature12335.

[25] Adelmann, J. A., Vetter, R. & Iber, D. The impact of cell size on morphogen gradient precision 150, dev201702. URL https://journals.biologists.com/dev/article/150/10/dev201702/310824/The-impact-of-cell-size-on-morphogen-gradient.

[26] Gόmez-Gálvez, P. et al. Scutoids are a geometrical solution to three-dimensional packing of epithelia 9, 2960. URL https://www.nature.com/articles/s41467-018-05376-1.

[27] Bocanegra-Moreno, L., Singh, A., Hannezo, E., Zagorski, M. & Kicheva, A. Cell cycle dynamics control fluidity of the developing mouse neuroepithelium URL https://www.nature.com/articles/s41567-023-01977-w.

[28] Guilak, F., Tedrow, J. R. & Burgkart, R. Viscoelastic properties of the cell nucleus 269, 781–786.

[29] Lammerding, J. in *Mechanics of the nucleus* 1 edn, (ed.Prakash, Y. S.) Comprehensive Physiology 783–807 (Wiley). URL https://onlinelibrary.wiley.com/doi/10.1002/cphy.c100038.

[30] Kalukula, Y., Stephens, A. D., Lammerding, J. & Gabriele, S. Mechanics and functional consequences of nuclear deformations 23, 583–602. URL https://www.nature.com/articles/s41580-022-00480-z.

[31] McGregor, A. L., Hsia, C.-R. & Lammerding, J. Squish and squeeze — the nucleus as a physical barrier during migration in confined environments 40, 32–40. URL https://linkinghub.elsevier.com/retrieve/pii/S0955067416300035.

[32] Cohen, R. et al. Mechanical forces drive ordered patterning of hair cells in the mammalian inner ear 11, 5137. URL https://www.nature.com/articles/s41467-020-18894-8.

[33] Kim, S. et al. A nuclear jamming transition in vertebrate organogenesis URL https://www.nature.com/articles/s41563-024-01972-3.

[34] Weigert, M., Schmidt, U., Haase, R., Sugawara, K. & Myers, G. Star-convex polyhedra for 3d object detection and segmentation in microscopy 1 (2019). URL http://arxiv.org/abs/1908.03636.

[35] Amini, R., Rocha-Martins, M. & Norden, C. Neuronal migration and lamination in the vertebrate retina 11, 1–16.

[36] Norden, C. A fish eye view: Retinal morphogenesis from optic cup to neuronal lamination 39, 175–196. URL https://www.annualreviews.org/doi/10.1146/annurev-cellbio-012023-013036.

[37] Hoon, M., Okawa, H., Della Santina, L. & Wong, R. O. Functional architecture of the retina: Development and disease 42, 44–84. URL https://linkinghub.elsevier.com/retrieve/pii/S135094621400038X.

[38] Buchsbaum, I. Y. & Cappello, S. Neuronal migration in the CNS during development and disease: insights from in vivo and in vitro models.

[39] Gleeson, J. G. & Walsh, C. A. Neuronal migration disorders: from genetic diseases to developmental mechanisms 23, 352–359.

[40] Cepko, C. Intrinsically different retinal progenitor cells produce specific types of progeny 15, 615–627. URL https://www.nature.com/articles/nrn3767.

[41] Levine, E. M. & Green, E. S. Cell-intrinsic regulators of proliferation in vertebrate retinal progenitors 15, 63–74. URL https://linkinghub.elsevier.com/retrieve/pii/S1084952103000648.

[42] Scott, G. D. & Kilgour, D. M. The density of random close packing of spheres 2, 863–866. URL https://iopscience.iop.org/article/10.1088/0022-3727/2/6/311.

[43] Berryman, J. G. Random close packing of hard spheres and disks. Phys. Rev. A 27, 1053–1061 (1983). URL https://link.aps.org/doi/10.1103/PhysRevA.27.1053.

[44] Torquato, S. & Stillinger, F. H. Jammed hard-particle packings: From kepler to bernal and beyond 82, 2633–2672. URL https://link.aps.org/doi/10.1103/RevModPhys.82.2633. Publisher: American Physical Society.

[45] Donev, A. et al. Improving the density of jammed disordered packings using ellipsoids 303, 990–993.

[46] Donev, A., Stillinger, F. H., Chaikin, P. M. & Torquato, S. Unusually dense crystal packings of ellipsoids 92, 255506. URL https://link.aps.org/doi/10.1103/PhysRevLett.92.255506.

[47] Nerli, E., Rocha-Martins, M. & Norden, C. Asymmetric neurogenic commitment of retinal progenitors involves notch through the endocytic pathway 9, 1–25.

[48] Maia-Gil, M. et al. Nuclear deformability facilitates apical nuclear migration in the developing zebrafish retina (2024). URL 10.1016/j.cub.2024.10.015. Publisher: Elsevier.

[49] Yanakieva, I., Erzberger, A., Mateǰcíc, M., Modes, C. D. & Norden, C. Cell and tissue morphology determine actin-dependent nuclear migration mechanisms in neuroepithelia 218, 3272–3289.

[50] Hong, E. & Brewster, R. N-cadherin is required for the polarized cell behaviors that drive neurulation in the zebrafish 133, 3895–3905. URL https://journals.biologists.com/dev/article/133/19/3895/52665/N-cadherin-is-required-for-the-polarized-cell.

[51] Ciruna, B., Jenny, A., Lee, D., Mlodzik, M. & Schier, A. F. Planar cell polarity signalling couples cell division and morphogenesis during neurulation 439, 220–224. URL https://www.nature.com/articles/nature04375.

[52] Hevia, C. F., Engel-Pizcueta, C., Udina, F. & Pujades, C. The neurogenic fate of the hindbrain boundaries relies on notch3-dependent asymmetric cell divisions 39, 110915. URL https://linkinghub.elsevier.com/retrieve/pii/S2211124722006921.

[53] Icha, J., Kunath, C., Rocha-Martins, M. & Norden, C. Independent modes of ganglion cell translocation ensure correct lamination of the zebrafish retina 215, 259–275.

[54] Salbreux, G., Barthel, L. K., Raymond, P. A. & Lubensky, D. K. Coupling mechanical deformations and planar cell polarity to create regular patterns in the zebrafish retina 8, e1002618. URL https://dx.plos.org/10.1371/journal.pcbi.1002618.

[55] Trushko, A. et al. Buckling of an epithelium growing under spherical confinement 54, 655–668.e6. URL https://linkinghub.elsevier.com/retrieve/pii/S1534580720305943.

[56] Stadler, J. A. et al. Histone deacetylase 1 is required for cell cycle exit and differentiation in the zebrafish retina 233, 883–889.

[57] Yamaguchi, M. et al. Histone deacetylase 1 regulates retinal neurogenesis in zebrafish by suppressing wnt ad notch signaling pathways 132, 3027–3043.

[58] Cunliffe, V. T. Histone deacetylase 1 is required to repress notch target gene expression during zebrafish neurogenesis and to maintain the production of motoneurones in response to hedgehog signalling 131, 2983–2995. URL https://journals.biologists.com/dev/article/131/12/2983/42249/Histone-deacetylase-1-is-required-to-repress-Notch.

[59] De Leeuw, N. F., Budhathoki, R., Russell, L. J., Loerke, D. & Blankenship, J. T. Nuclei as mechanical bumpers during epithelial remodeling 223, e202405078. URL https://rupress.org/jcb/article/223/12/e202405078/276995/Nuclei-as-mechanical-bumpers-during-epithelial.

[60] Hannezo, E., Prost, J. & Joanny, J.-F. Theory of epithelial sheet morphology in three dimensions 111, 27–32. URL https://pnas.org/doi/full/10.1073/pnas.1312076111.

[61] Drasdo, D. Buckling instabilities of one-layered growing tissues 84, 4244–4247. URL https://link.aps.org/doi/10.1103/PhysRevLett.84.4244.

[62] Franze, K. et al. Müller cells are living optical fibers in the vertebrate retina 104, 8287–8292. URL https://pnas.org/doi/full/10.1073/pnas.0611180104.

[63] Reichenbach, A., Agte, S., Francke, M. & Franze, K. How light traverses the inverted vertebrate retina: No flaw of nature 5, 93–100. URL http://link.springer.com/10.1007/s13295-014-0054-8.

[64] Solovei, I. et al. Nuclear architecture of rod photoreceptor cells adapts to vision in mammalian evolution 137, 356–368. URL https://linkinghub.elsevier.com/retrieve/pii/S0092867409001378.

[65] Kreysing, M., Boyde, L., Guck, J. & Chalut, K. J. Physical insight into light scattering by photoreceptor cell nuclei 35, 2639. URL https://opg.optica.org/abstract.cfm?URI=ol-35-15-2639.

[66] Kimmel, C. B., Ballard, W. W., Kimmel, S. R., Ullmann, B. & Schilling, T. F. Stages of embryonic development of the zebrafish 203, 253–310.

[67] Geldmacher-Voss, B., Reugels, A. M., Pauls, S. & Campos-Ortega, J. A. A 90° rotation of the mitotic spindle changes the orientation of mitoses of zebrafish neuroepithelial cells 130, 3767–3780. URL https://journals.biologists.com/dev/article/130/16/3767/52119/A-90-rotation-of-the-mitotic-spindle-changes-the.

68. Dzafic, E., Strzyz, P. J., Wilsch-Bräuninger, M. & Norden, C. Centriole amplification in zebrafish affects proliferation and survival but not differentiation of neural progenitor cells 13, 168–182. URL https://linkinghub.elsevier.com/retrieve/pii/S2211124715009596.

[69] Zolessi, F. R., Poggi, L., Wilkinson, C. J., Chien, C.-B. & Harris, W. A. Polarization and orientation of retinal ganglion cells in vivo 1, 2. URL https://neuraldevelopment.biomedcentral.com/articles/10.1186/1749-8104-1-2.

[70] Soans, K. G. et al. Collective cell migration during optic cup formation features changing cell-matrix interactions linked to matrix topology 32, 4817–4831.e9. URL https://linkinghub.elsevier.com/retrieve/pii/S0960982222015032.

[71] Schindelin, J. et al. Fiji: an open-source platform for biological-image analysis 9, 676–682. URL https://www.nature.com/articles/nmeth.2019.

[72] Machado, S., Mercier, V. & Chiaruttini, N. LimeSeg: a coarse-grained lipid membrane simulation for 3d image segmentation 20, 2. URL https://bmcbioinformatics.biomedcentral.com/articles/10.1186/s12859-018-2471-0.

[73] Sofroniew, N. et al. napari: a multi-dimensional image viewer for python. URL https://zenodo.org/doi/10.5281/zenodo.3555620.

[74] Archit, A. et al. Segment anything for microscopy. URL http://biorxiv.org/lookup/doi/10.1101/2023.08.21.554208.

## References

[1] A. Trushko, I. Di Meglio, A. Merzouki, C. Blanch-Mercader, et al., Dev. Cell 54(5) 655–668 (2020).

[2] G. Napoli and S. Turzi, Proc. R. Soc. A 471 2183 (2015).

[3] A.L. McGregor, C.-R. Hsia, and J. Lammerding, Curr. Opin. Cell Biol. 40 32–40 (2016).

[4] Y. Kalukula, A.D. Stephens, J. Lammerding, and S. Gabriele, Nat. Rev. Mol. Cell Biol. 23 583–602 (2022).

[5] A.E.H. Love, A Treatise on the Mathematical Theory of Elasticity, 4th Ed., Cambridge University Press (1927).

